# WITER: A powerful method for the estimation of cancer-driver genes using a weighted iterative regression accurately modelling background mutation rate

**DOI:** 10.1101/437061

**Authors:** Lin Jiang, Jingjing Zheng, Johnny Sheung Him Kwan, Sheng Dai, Cong Li, Mulin Jun Li, Bolan Yu, Ka Fai TO, Pak Chung Sham, Yonghong Zhu, Miaoxin Li

## Abstract

Genomic identification of driver mutations and genes in cancer cells are critical for precision medicine. Due to difficulty in modeling distribution of background mutations, existing statistical methods are often underpowered to discriminate driver genes from passenger genes. Here we propose a novel statistical approach, weighted iterative zero-truncated negative-binomial regression (WITER), to detect cancer-driver genes showing an excess of somatic mutations. By solving the problem of inaccurately modeling background mutations, this approach works even in small or moderate samples. Compared to alternative methods, it detected more significant and cancer-consensus genes in all tested cancers. Applying this approach, we estimated 178 driver genes in 26 different cancers types. *In silico* validation confirmed 90.5% of predicted genes as likely known drivers and 7 genes unique for individual cancers as likely new drivers. The technical advances of WITER enable the detection of driver genes in TCGA datasets as small as 30 subjects, rescuing more genes missed by alternative tools.

## Introduction

It is well known that genomic aberration in somatic cells makes important contribution to development of cancers(1). Mutations that confer selective growth advantage to cancer cells are called as cancer-drivers (2) (3); a gene harboring driver-mutations is defined as a cancer-driver gene. It has been established, for example, that mutations in the two famous driver genes TP53 and PIK3CA contribute to many types of cancer (4). However, cancers are known to be highly heterogeneous(5) and many driver genes of most cancers remain to be identified. A full landscape of driver-genes, which is fundamental for early diagnosis, identification of effective drug targets, and precise treatments of cancers (2), remains unavailable for most cancers.

There are generally two existing strategies to detect cancer driver genes with somatic mutations, background mutation rate (BMR) and ratiometric. The BMR-based methods evaluate whether a gene has more somatic mutations than expected; examples include MutSigCV (6) and MuSiC(7). The expected number of mutations is estimated by multiple predictors including base context, gene size and other variables of genes. The ratiometric-based methods detect cancer-driver genes according to the composition of mutation types normalized by the total number of mutations in a gene. For instance, the ratiometric 20/20 rule assessed the proportion of inactivating mutations (including synonymous mutations) and missense mutations(3). Oncodrive-fm(8) and OncodriveFML(9) integrated mutations’ functional impact into the evaluation. OncodriveCLUST considered the positional clustering of mutation patterns(10). Recently a method 20/20 plus (11) extended the ratiometric idea in the 20/20 rule and integrated 18 additional features of positive selection to predict cancer-driver genes by a machine learning approach. It also generated statistical p-values of the prediction scores by Monte Carlo simulations.

Although the general principles of both strategies are simple, three key technical issues remain unsolved. First, the background mutation rates of genes are not accurately modelled generally. A recent study (11) found that the statistical p-values produced by existing cancer-driver gene methods did not follow uniform distribution, implying the underfitting of background mutations. Although simulation or permutation can be used correct the p-values, adequate fitting of background genes is critical for accurate discrimination of true driver genes from noise background genes. Second, existing statistical tests are generally underpowered to detect driver-genes with small or moderate effect size. This issue will become more severe when the sample is small. This may be a reason why some supervised approaches integrating common gene features beyond collected samples were also proposed(11). However, given the high heterogeneity in cancers (6), adding more common features may not work for unique driver genes of a cancer. The trained model for known driver genes may have limited power for detecting new driver genes. Lastly, the predicted cancer-driver genes by different tools do not generally agree with each other(2). It is often laborious and subjectively biased to combine their results. Therefore, more powerful methods are pressingly needed for unraveling a full spectrum of cancer-driver genes.

Here, we describe a new statistical method, weighted iterative zero-truncated negative-binomial regression (WITER), to detect cancer-driver genes by somatic mutations [including single nucleotide variants (SNVs) and short insertions and deletions InDels]. This approach belongs to the unsupervised category and therefore does not suffer from training bias. The method has a unique three-tier structure to ensure accurate fitting of the somatic mutations of background genes even in small samples. We then systematically compare its performance with alternative methods in 11 cancers. A comprehensive landscape of driver-genes is constructed by WITER and analyzed to investigate the common and unique insights across 26 cancers.

## Methods and Materials

### The unified statistical framework for detecting cancer-driver genes by somatic mutations in cancers

We propose a unified statistical framework, WITER, for detecting cancer-driver genes by somatic mutations in cancers. The main input is somatic mutations (including SNVs and InDels) in samples from cancer patients. The output is a table of p-values for excess of somatic mutations at individual genes; a significant p-value suggests a driver gene that has more somatic mutations because such mutations confer selective growth advantages to cancer cells. The unified statistical framework has a three-tier structure to examine driver genes by using somatic mutations in cancer cells (See the diagram in Figure 1 and the following details). These methods work from different angles to improve the modelling of background mutations in passenger genes for a more powerful evaluation of driver genes. In theory, the framework and the model are independent of types of somatic variants to be tested. However, the present paper focuses on non-synonymous and splicing variants because of abundant validation data and resource data in public domains. The approach and auxiliary functions have been implemented into a user-friendly software tool which is publicly available at http://grass.cgs.hku.hk/limx/witer.

**Figure 1:**
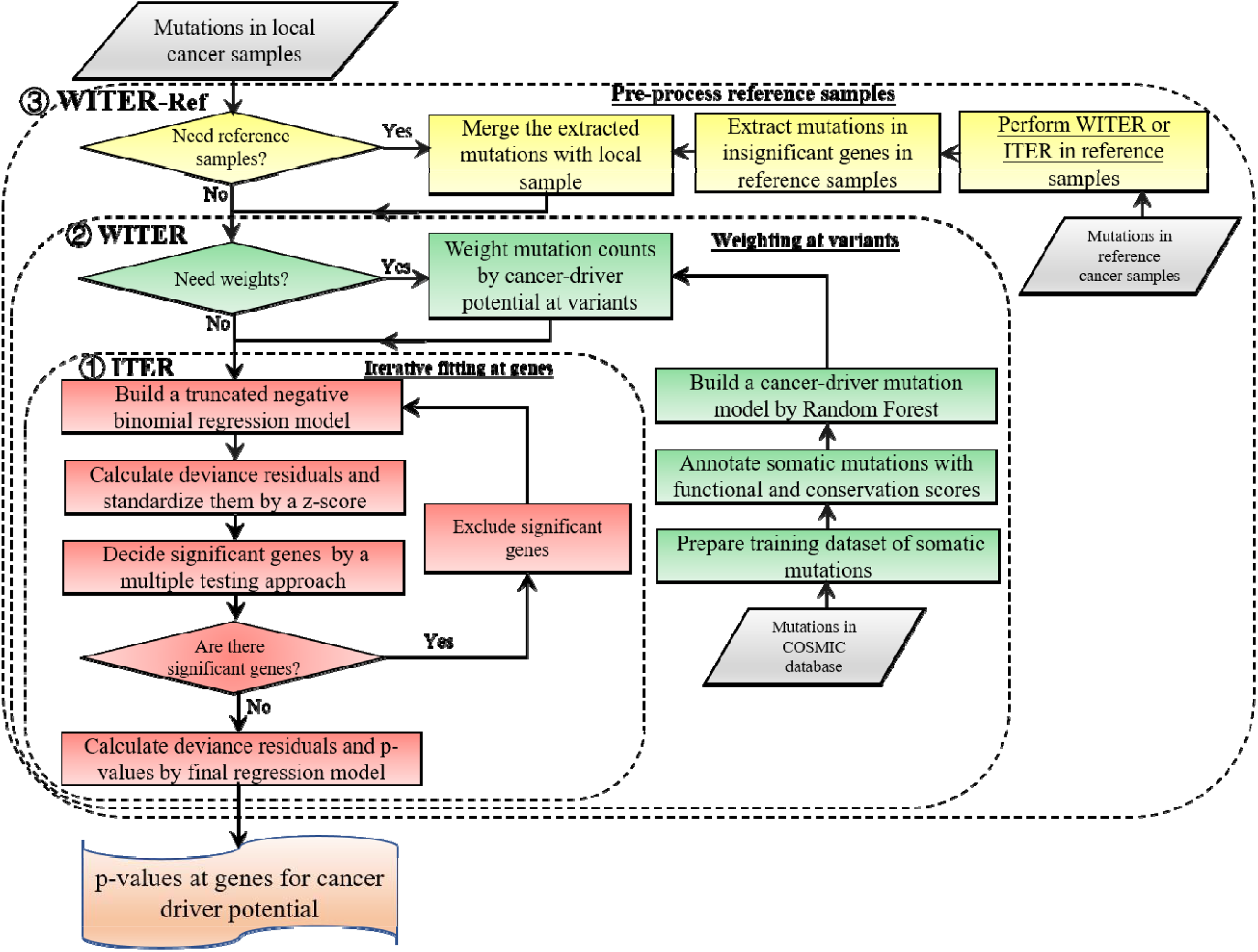
The diagram of the statistical framework for detecting cancer-driver genes. This framework includes three tiers denoted by the dashed rectangles. The first tier is an iterative zero-truncated negative-binomial regression (ITER). The second tier is a weighted iterative zero-truncated negative-binomial regression (WITER). The third tier is the integration of reference samples. The unique components of each tier are marked by different colors. The major inputs are somatic mutations in different cancer patients. The outputs are *p*-values for excess of somatic mutations of individual gene in the cancer samples.

#### Tier I: An iterative zero-truncated negative-binomial regression to model background somatic mutations

We proposed an approach, iterative zero-truncated negative-binomial regression (ITER), to estimate somatic mutation counts of each gene on the genome. The difference between the observed mutation counts and the estimated counts of a gene measures the excess of somatic mutations at a gene in a cancer. The assumption is that a gene with significant excess of somatic mutations may confer selective growth advantage in cancer as a driver gene(6). Denote the mutant allele counts at a variant *j* in a background gene *i* as *c*_*i,j*_ and the total alleles of *m*_*i*_ variants in this gene is, *y*_*i*_. We assume *y*_*i*_ follows a negative binomial (NB) distribution:

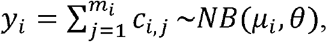

where *μ*_*i*_ is the expected number of mutations and *θ* is a dispersion parameter. The probability mass function (PMF) is 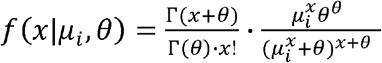, where Γ( ) is the gamma function and *x*=0,1,2, ….

As somatic mutation is a rare event, many genes have no somatic mutations in a sample of typical size. While the negative binomial model includes a probability mass at *x*=0, this is often much smaller than the proportion of genes without somatic mutations in real data. This inflation of zeros makes it very difficult to fit the negative binomial distribution to the counts of somatic mutations. Therefore, we proposed to use a zero-truncated negative binomial (ZTNB) distribution to model the mutant allele counts of background gene *i*. The PMF of ZTNB is:

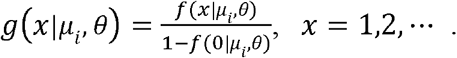

Based on the ZTNB, we constructed a generalized linear regression model to estimate mutant allele of non-synonymous or splicing variants in a gene *i* by 6 covariables:

*η* = log(*η*_*i*_) = *β*_0_ + *β*_1_ × [*x*_1_, length of genomic regions of the variant types] + *β*_2_ × [*x*_2_,…] + · · · + *β*_*m*_ × [*x*_*m*_,…], where log (*η*_*i*_) is the link function and the *β*s are the regression coefficients.

The key idea is the ZTNB distribution for somatic mutations in background or passenger genes. We showed the ZTNB model fitted the mutations counts better than other alternative models in the Results section (Table 3). Besides the length of genomic regions, the predictors in the regression model are flexible and depend on the available resources. Several studies showed somatic mutation rates tend to be higher in genes with low expression levels, repressed chromatin, DNA modification, and late replication times in cancer cells (6,12-14). Therefore, the four types of predictors are adopted in the prediction models. Besides, copy numbers variations (CNVs) occur in cancer cells frequently (15). It may be necessary to recalibrate the background mutation rate by the CNVs of background genes although the CNVs of driver genes may also contribute to cancers (15). In addition, assuming the synonymous and non-synonymous mutations in the same passenger genes have similar mutation burden, we also consider the number of synonymous mutations in local cancer samples as a predictor. The last predictor we used in the present paper is gene’s constraint scores for non-synonymous mutations in natural populations (16), which assumes a gene having higher mutation potential in germline cells tends to have higher mutation potential in somatic cells as well. There are in total 10 predictors in the present paper (See details in Table 6). This model is also open for other types covariables as long as they can improve the prediction accuracy. Statistically insignificant predictors will have little contribution to the estimation.

The coefficients of the predictors can be estimated by maximum likelihood with a quasi-Newton method. With the estimated coefficients, the deviance residues are calculated and standardized as *é*_*i*_ (See the detailed methods in the supplementary notes). A large *é*_*i*_ means the observed number of somatic mutations is larger than the expected number of mutations under null hypothesis. The corresponding p-value is then approximated by,

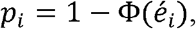

where *Φ*(*x*) is the cumulative distribution function of the standard normal distribution. We demonstrate the p-values follow uniform distribution in real data analysis [Figure 2, 3a and S1].

**Figure 2:**
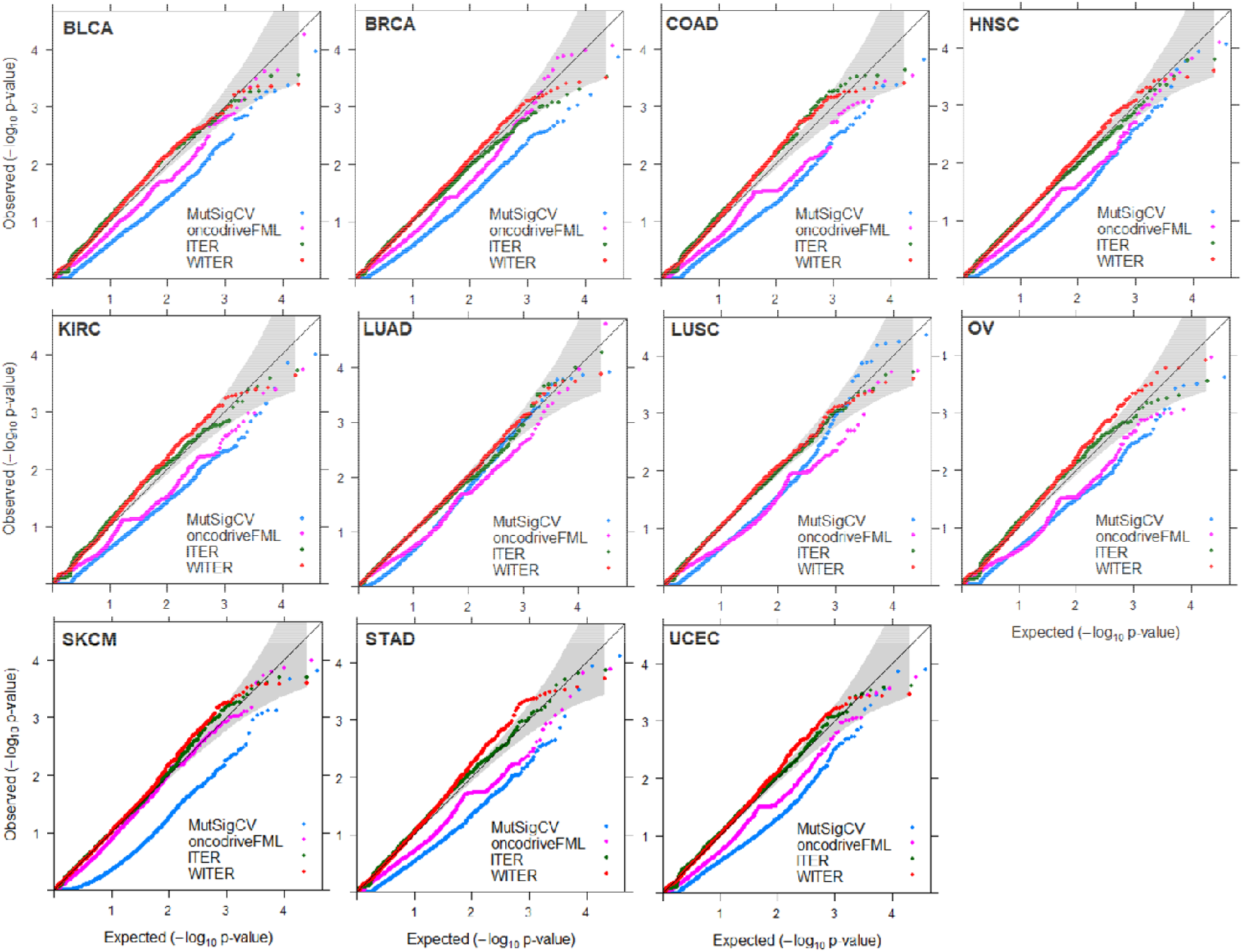
QQ plot of background gene p-values produced by 4 methods in 11 cancers. The p-values less than a cutoff according to FWER 0.05 were excluded. Among the 34 collected cancers, 11 cancers have 25,000 variants with somatic mutations in the data sets and are used for the comparison.

**Figure 3:**
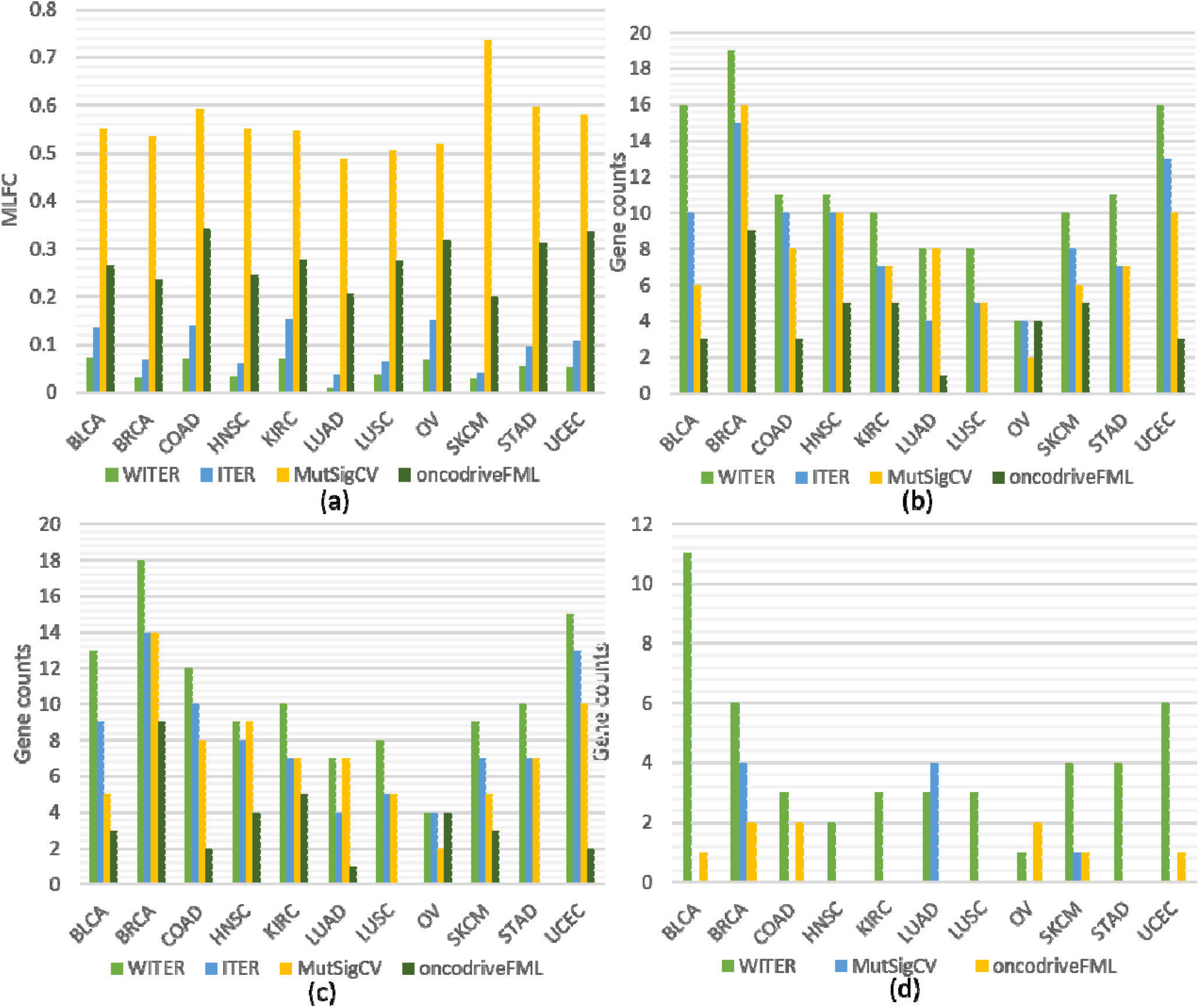
Performance comparison of different methods for detecting cancer driver mutation in 11 cancers. a: The MLFC of 4 methods; b: the number of significant genes; c: cancer consensus significant genes; d: the number of unique significant genes. The p-values less than a cutoff according to FWER 0.05 are excluded. The full names of cancers are in Table S1.

Note the above ZTNB model is used to estimate somatic mutations among passenger genes. The mutation counts in drive genes may harm the estimation. In order to reduce distortion of driver genes, we further propose to perform the regression under an iterative procedure:

Step 1: fit ZTNB and calculate p-vales for all genes.

Step 2: exclude significant genes by a loose p-value cutoff corresponding to false discovery rate (FDR)≤0.1.

Step 3: fit ZTNB and calculate p-vales for the retained genes.

Step 4: repeat Step 2 and 3 until there is no extra significant genes according to the same p-value cutoff.

The fitted ZTNB model in the last iteration is closest to the null hypothesis model and is then used to re-calculate deviance residuals and p-values of all genes (including the ones excluded during iteration).

#### Tier II: A weighting scheme to prioritize variants of high somatic mutation potential in cancer samples

We further extend ITER to a WITER, which integrates prior weights at variants to boost power. Assume a variant *j* of gene *i* has a score, *s*_*i,j*_ ∈ [0,1], implying its cancer driver potential. We bin *s*_*i,j*_ as an integer score, *w*_*i,i*_, by the ceiling function of *s*_*i,j*_/0.3, i.e., *w*_*i,i*_ = ⌈ *s*_*i,j*_/0.3 ⌉. The integer score is then used as prior weights for the variant. The ITER is a special case of WITER when *w*_*i,j*_ = 1 for all variants. The weighted mutation allele count is:

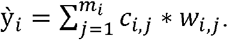

We also assume the weighted counts 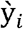 follow a negative binomial (NB) distribution:

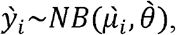

where 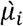 is the expected weighted count of mutations and 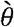 is a dispersion parameter of the NB distribution. After replacement of original counts (*y*_*i*_) with weighted counts 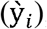, the same iterative ZTNB regression procedure is carried out to test whether a gene has excess of weighted mutant alleles.

In the present study, we built a model to predict high-frequency cancer driver potential to use as prior weights, by the random forest trained in a large cancer somatic mutation database, COSMIC (V83). (See details in the supplementary notes). One can also use other methods to produce the prior weights.

#### Tier III: A schedule of integrate independent reference samples to stabilize the regression model for small samples

When the sample size is small, it is difficult to build a stable regression model. Note that the key idea of ITER and WITER is to build a prediction model for background passenger genes. When the mutation rates of passenger genes of two cancers are similar, it may be workable to integrate background genes of one cancer into another cancer. We proposed a reference sample strategy for construct a stable ITER or WITER model in small samples. This is carried out at two stages.

i. The above ITER or WITER is used to produce p-values for excess of somatic mutations at genes in a reference sample which have enough variants. Genes with p-values less than a very loose cutoff, say FDR 0.3, are then excluded.
ii. The somatic mutations of retained genes are integrated with the local small sample and input into ITER or WITER to build a new model. The excess of somatic mutations and corresponding p-values at genes are calculated based on the new model.

### Curation of cancer-specific predictors of somatic mutations

We collected 4 types of cancer-specific predictors for somatic mutations, copy number variation (CNV), gene expression, DNA methylation, and chromatin accessibility by ATAC-Seq. All data were produced from TCGA cohorts and the preprocessed data were downloaded from https://xenabrowser.net. The pan-CNVs mapped onto genes were downloaded by the link, https://tcga.xenahubs.net/download/TCGA.PANCAN.sampleMap/Gistic2_CopyNumber_Gistic2_all_thresholded.by_genes.gz. We then wrote a program to group subjects according to cancer types. The mean CNV of a gene in all subjects of the same cancer type was used as the CNV of the gene at the cancer type. The batch-effect-normalized gene expression was downloaded from https://pancanatlas.xenahubs.net/download/EB++AdjustPANCAN_IlluminaHiSeq_RNASeqV2.geneExp.xena.gz. Similar to CNV, the mean expression of a gene at a cancer type was also used as a predictor in the regression model. The DNA methylation assayed by illuminaMethyl450 was downloaded from https://pancanatlas.xenahubs.net/download/jhu-usc.edu_PANCAN_HumanMethylation450.betaValue_whitelisted.tsv.synapse_download_5096262.xena.gz. We wrote a program to map the methylation regions and beta-values onto exons of genes according to the gene models defined in RefGene and GENCODE. The overall DNA methylation proportion in all exons of a gene was used as the DNA methylation level of the gene of a subject. A gene’s averaged methylation level in all subjects of the same cancer types was used as a cancer specific predictor in the regression model. The peak and region of ATAC-seq for chromatin accessibility were downloaded from https://atacseq.xenahubs.net/download/TCGA_ATAC_peak_Log2Counts_dedup_sample.

Similar to the methylation level, the ATAC-seq peaks and regions were mapped onto exons of each gene. The overall ATAC-seq signals (coverage multiplying peaks) at all exons of each gene were calculated for each subject. A gene’s averaged ATAC-seq signal in all subjects of the same cancer types was used as a cancer specific predictor in the regression model.

### Performance comparison with alternative tools

There have been multiple tools for detecting cancer-driver genes(2). According to an evaluation study(11), three tools (MutSigCV(6), OncodriveFML(9) and 20/20plus(11)) having relatively better performance were chosen for comparisons in the present study. We compared their p-value distributions and number of significant genes to ITER and WITER. As MutSigCV and OncodriveFML were also developed under an unsupervised strategy, we chose them as the main comparison targets. The 20/20plus belongs to a supervised strategy which may be more suitable for known cancer driver genes. To be fair, we only used it as supplementary comparison. The details of the usage of the alternative methods are described in supplementary notes.

### Evaluation metrics in the performance comparison

We adopted four evaluation metrics for performance comparison, number of significant genes predicted, overlap with Cancer Gene Census (CGC) (17), observed vs. theoretical *p* values, and percentage of CGC genes in significant genes. The former 3 were also major metrics in an evaluation framework of cancer driver gene prediction method(11). The CGC dataset contained 699 manually curated cancer genes. The departure of p-values from uniform distribution was measured by the mean absolute log2 fold change (MLFC) (11). A valid statistical test should lead to a MLFC close to zero in background (or passage) genes. We also used the distribution of Quantile-Quantile (QQ) plot to examine the distribution of p-values, particularly that of the small p-values. The Bonferroni correction for family-wise error rate (FWER) 0.05 was used to report significant genes.

### Dataset of somatic mutations

We partitioned a curated full somatic mutation dataset by Tokheima and colleagues (11) into 34 sub-datasets according to the cancer types (See Table S1). Eleven cancer types contained 2,800 or more variants (See the full list in Table S1). These cancers samples were called relatively larger dataset throughout the paper and used for the method comparison. Their sample sizes ranged from 142 to 1093. The ratios of variant number to sample size in the 11 cancers ranged from 50 to 327. The remaining 23 cancers with a smaller number of variants were only used in the application analysis. The names, variant number and sample sizes of all cancers can be seen in Table S1.

### In silico validation by PubMed search

We used PubMed search function to coarsely validate the implication of detected significant genes the corresponding cancer. The underlying assumption is that the papers co-mentioning the gene and the cancer name in the title or abstract are likely to implicate the relatedness between the gene and the cancer. The more hit papers, the more likely the gene is related to the cancer. This is a quick *in-silico* validation although it may be rough. We employed the web application programming interfaces (APIs) of PubMed to execute the search. The search link was, http://eutils.ncbi.nlm.nih.gov/entrez/eutils/esearch.fcgi?db=pubmed&term=“DiseaseNames(inlcudinghomonymies)”[tiab]%29+AND+“GeneSymbol (including RefSeq mRNA IDs)” [tiab]. The search terms of each cancer types are in Table S1. The search responded PubMed IDs and relevant data of the papers, if available.

## Results

### Features correlated with somatic mutations of genes in cancer samples

We first investigated association of the 10 individually explanatory features with somatic mutation counts under the ZTNB regression in 11 cancers(See coefficients and p-values in Table 2). While confirming previous finding that somatic mutation rates tend to be higher in genes with low expression levels, repressed chromatin, and late replication times in most cancers (6,12-14), there are also four interesting patterns. First, for gene expression and chromatin state, it seems cancer non-specific features are more significantly correlated with the somatic mutations than the cancer specific features in most cancers. For example, the averaged expression level in cancer cell lines has much more significant p-values in 10 out of the 11 cancers than the gene expression level in the matched cancer tissues. The chromatin state assayed by HiC from the K562 cell line is much more significant than that assayed by cancer-matched ATACSeq in 10 out of the 11 cancers. The absolute coefficient values of the former are also larger than that of the latter in the 10 cancers. Note all the feature values are standardized to make the coefficients comparable. Second, the significance level of most features varies from cancers to cancers. This is particularly true for three features, CNV, methylation and constraint score. For instance, CNV has a very significant p-value, 9.71E-20, in LUAD while its p-value is only 0.16 in BLCA. For methylation, its smallest p-value occurs in KIRC (p=1.57E-9) while it has large p-values (>0.3) in 4 cancers. Third, besides the varied significance level, some features’ association directions are also different in different cancer types. The constraint score has positive significant coefficients in BRCA, KIRC and UCEC while it has negative significant coefficients in LUAD, LUSC and SKCM. While genes with higher CNV tend to have more somatic mutations in four cancers (HNSC, LUAD and LUSC), the tendency gets reversed in KIRC. The underlying mechanisms are unclear. Fourth, five cancer non-specific features (exon length, number of synonymous mutations, replication timing and cell line expression levels and HiC) have extremely significant p-values for all cancers. These features dominate the prediction performance in the ZTNB model. The advantage in cancer non-specificity makes the methods more flexible and feasible in practice.

**Table 1.**
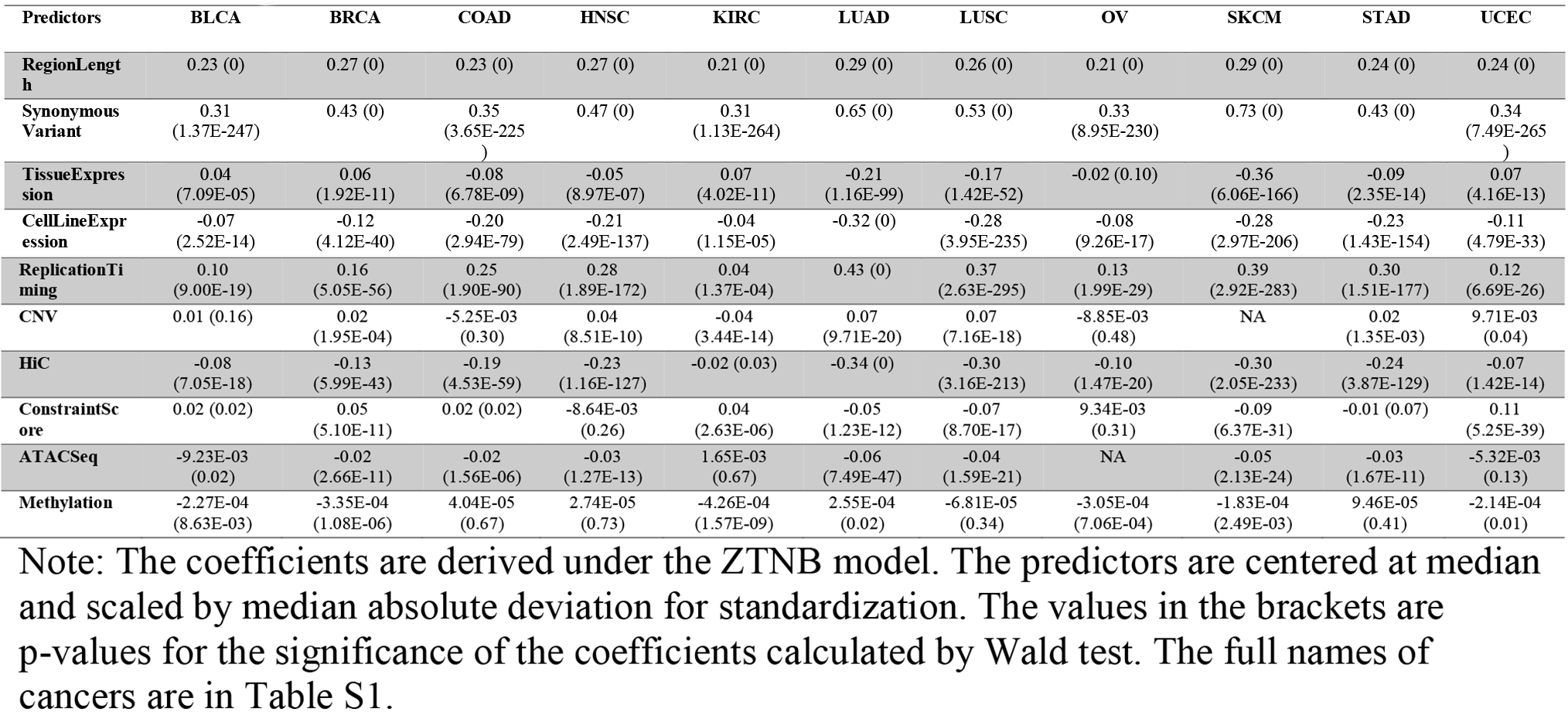
The significance level of covariates in 11 cancer datasets

**Table 2:** Akaike information criterion (AIC) of the various regression models

|  | Poisson | Zt- Poisson | Negative Binomial | Zt-Negative Binomial |
| --- | --- | --- | --- | --- |
| BLCA | 50439.28 | 49991.30 | 42766.17 | 40965.04 |
| BRCA | 93502.18 | 93343.70 | 60491.40 | 58701.48 |
| COAD | 55069.59 | 54675.87 | 40495.83 | 38459.15 |
| HNSC | 91759.62 | 91608.29 | 60737.90 | 59131.56 |
| KIRC | 49696.06 | 49152.82 | 38404.63 | 36484.36 |
| LUAD | 140194.62 | 140168.95 | 79081.09 | 78048.05 |
| LUSC | 79647.23 | 79409.43 | 55650.58 | 54129.60 |
| OV | 53288.73 | 52812.71 | 40753.37 | 38623.84 |
| SKCM | 126186.28 | 126133.21 | 70968.39 | 69622.20 |
| STAD | 63094.87 | 62754.31 | 48865.94 | 47114.14 |
| UCEC | 66024.20 | 65665.04 | 48584.07 | 46526.46 |

Note: The glm() function in R was used to fit the generalized linear model (GLM) of Poisson distribution. The glm.nb() function in the R package of MASS was used to fit the GLM of Negative Binomial distribution. The other two models were fitted by the R package of countreg. The full names of cancers are in Table S1.

### Distributions of *p*-values and fitness of the ZTNB model for background mutation genes

We then investigated the type 1 errors of WITER and ITER according to the distribution of p-values. When the overall divergence from uniform distribution are measured as the MLFC for insignificant p-values (FWER>0.05) (11), the WITER has small MLFC (<0.1) in all the tested 11 cancers (Figure 3a). The smaller MLFC, the closer to uniform distribution. We also choose two alternative approaches which achieved the best performance among 7 widely-used unsupervised tools(11) for the comparison, MusigCV (6) and OncodriveFML (9). Among the four compared methods, ITER has the second smallest MLFC. MusigCV (6) has the largest MLFC values, in which most values are over than 0.5. In terms of the MLFC, the OncodriveFML (9) is better than MusigCV but worse than ITER. As shown in the QQ plot (Figure 2), the main problem in MusigCV and OncodriveFML is their deflated p-values although the deflation at small p-values genes is mitigated. Therefore, the proposed ZTNB model addressed the problem of invalid uniform distribution of *p*-values, which is also a common challenge in almost all existing approaches (11).

We also compared the goodness-of-fit of the ZTNB model with three widely-used alternative models for somatic mutation counts of genes (Table 2). It turns out the ZTNB model has the smallest Akaike information criterion (AIC) values in all the 11 cancers, suggesting the best goodness of fit among the four models (Table 2). The negative binomial distribution is the second best model although its AIC values are over 1000 larger than that of the ZTNB model in all cancers. The Poisson, either in the original version or the zero-truncated version, fit data poorly. Their AIC values are much larger than that of the ZTNB. The best goodness-of-fit reaffirms that ZTNB is a suitable model for the counts of somatic mutations in background genes. The well-fitted ZTNB model leads to more accurate residues for evaluating the relative excess of mutations in driver genes.

### WITER detected more significant genes in 11 cancers

We compared the number of significant genes detected by the 4 unsupervised approaches (MutSigCV, OncodriveFML, ITER and WITER) in the 11 cancer datasets. Instead of following the conventional “pancancer” (all cancers) evaluation strategy (11), we made the comparison for individual cancers, a more challenging scenario because of smaller sample sizes. The significant genes are determined according to Bonferroni correction for FWER 0.05. Among the 4 approaches, WITER estimates the largest number of significant genes in all cancers (Figure 3b). ITER estimates the second largest number of significant genes in 9 out of the 11 cancers. MutSigCV can be ranked at the third place according to the number of significant genes. The OncodriveFML detects the minimal number of significant genes in 10 out of the 11 cancers although it also integrates function prediction score CADD (18). Given the correct type 1 errors, the more significant genes by WITER than ITER suggests the prior functional weights at variants have great potential to improve the statistical power. Note that all the subjects in the testing cancer datasets were excluded from the COSMIC database to avoid circulating issues when building the prior weights for WITER.

A critical follow-up question is that whether the increased number of significant genes by WITER or ITER are true driver genes. As there are almost few true answers in the real data, we adopt the Cancer Gene Census (CGC) list (17) to partly address this question. Among the significant genes, WITER always leads to the largest number of CGC genes in all the 11 cancers (Figure 3c). It detects at least 3 and 6 more CGC genes in 8 different cancers than MutSigCV and OncodriveFML respectively. ITER is still the second-best method according to the number of CGC genes. It detects more CGC genes than MutSigCV in 5 cancers and has equal number of CGC genes with MutSigCV in 4 cancers. Compared to OncodriveFML, ITER detects more CGC genes in 10 cancers. Moreover, we also check the percentage of CGC genes among significant genes, which might imply the false positive rates to some extent. The percentage varies from cancers to cancers. Compared to MutSigCV, WITER has higher or equal percentage in 7 out of the 11 cancers (See details in Table 3). The averaged percentages of WITER and MutSigCV in the cancers are comparable, 91.2% and 93.8%. ITER has the highest or equal percentage in 8 out of the 11 cancers and achieves a slightly better averaged percentage, 94.1%. Although OncodriveFML has the smallest number of significant genes, it has the smallest averaged percentage, 85.9%. This is because it has low percentages of 60%, 66.6% and 66.6% in three cancers, SKCM, UCEC and COAD(See details in Table 3).

**Table 3:** The percentage of the cancer consensus gene in the significant genes by different methods

|  | WITER | ITER | MutSigCV | OncodriveFML |
| --- | --- | --- | --- | --- |
| BLCA | 76.5 | 81.8 | 83.3 | 100 |
| BRCA | 94.7 | 100 | 87.5 | 100 |
| COAD | 100 | 100 | 100 | 66.7 |
| HNSC | 75 | 72.7 | 90 | 80 |
| KIRC | 100 | 100 | 100 | 100 |
| LUAD | 87.5 | 100 | 87.5 | 100 |
| LUSC | 100 | 100 | 100 | - |
| OV | 100 | 100 | 100 | 100 |
| SKCM | 90 | 87.5 | 83.3 | 60 |
| STAD | 90.9 | 100 | 100 | - |
| UCEC | 88.2 | 92.9 | 100 | 66.7 |
| **Average** | 91.2 | 94.1 | 93.8 | 85.9 |

Note: The full names of cancers are in Table S1.

**Table 4:** Rescued genes in the half-sample by different tools

| Tested tool<sup>a</sup> | RandomS<br>ampleID | SubSample | Missed | Rescued by |  |  |
| --- | --- | --- | --- | --- | --- | --- |
|  |  | Sig.Genes<sup>b</sup> | Sig.Genes | MutSigC<br>V | oncodriveFM<br>L | WITER |
| **MutSigCV**<br>**(16)** | 1 | 10 | 6 | -c | 0 | 2 |
|  | 2 | 8 | 8 | - | 0 | 3 |
|  | 3 | 10 | 6 | - | 0 | 3 |
|  | 4 | 7 | 9 | - | 1 | 3 |
|  | 5 | 8 | 8 | - | 2 | 4 |
|  | 6 | 8 | 8 | - | 0 | 4 |
| **OncodriveFM**<br>**L (9)** | 1 | 5 | 4 | 0 | - | 0 |
|  | 2 | 5 | 4 | 0 | - | 2 |
|  | 3 | 6 | 3 | 0 | - | 1 |
|  | 4 | 6 | 3 | 0 | - | 1 |
|  | 5 | 5 | 4 | 0 | - | 1 |
|  | 6 | 6 | 3 | 0 | - | 2 |
| **WITER (19)** | 1 | 14 | 5 | 0 | 0 | - |
|  | 2 | 16 | 3 | 0 | 0 | - |
|  | 3 | 14 | 5 | 0 | 1 | - |
|  | 4 | 14 | 5 | 0 | 0 | - |
|  | 5 | 14 | 5 | 0 | 0 | - |
|  | 6 | 17 | 2 | 0 | 0 | - |

Note: This experiment is performed in the BRCA dataset. a: The number of significant genes detected by a tool in the full dataset. b: The number of significant genes detected by a tool in the randomly extracted half samples. c: This item is inapplicable.

Note that the significant genes beyond CGC list are not necessarily spurious driver genes although a high percentage and number of CGC genes is a strong sign of higher power. Take two non-CGC genes for examples. The AJUBA gene (*p*=2.7E-8 in HNSC) is involved in the regulation of NOTCH/CTNNB1 signaling and is an important driver gene of HNSC(19) (20). TLR4 (*p*=3.7E-6 in STAD) is an important member of Toll-like receptor (TLR) pathway and mutations in the gene may disrupt innate immune signaling and promote a microenvironment that favors tumorigenesis (21) and it was associated with gastric cancer in independent samples (22).

### Uniquely significant genes by individual approaches

We also investigated genes only significant in one of the compared tools. WITER detects the largest number of uniquely significant genes (FWER≤0.05) in the 11 cancer types, which are insignificant and would be ignored by MutSigCV and OncodriveFML (Figure 3d). WITER detects in total 46 uniquely significant genes in all 11 cancers, among which 37 (80.4%) genes are in the CGC list, (enrichment *p*=1.8E^-45^, by hypergeometric distribution test in 19198 protein coding genes). The BLCA has the largest number, 11, of uniquely significant genes by WITER among which 8 genes are CGC genes. Some genes are well-known driver genes for BLCA, e.g., ERBB3(23). ERBB3 has 17 non-synonymous somatic mutant alleles in the BLCA samples. WITER calculates a p-value 7.2E-8 at this gene. The p-values by MutSigCV and OncodriveFML are 0.0016 and 0.25 respectively. MutSigCV does not detect any unique significant genes in 8 cancers. It detects in total 9 uniquely significant genes in three cancers among which 5 (=55.6%) genes are in the CGC list. For example, a non-CGC gene, FBN2, has 76 non-synonymous or splicing mutant alleles in the LUAD samples and MutSigCV produces a p-value 4.28E-07. However, it has a 9.1 kb coding region, 10 synonymous mutant alleles and late replication timing and low expression in LUAD tissues, WITER produces an insignificant p-value 0.16 for its excess of the adjusted non-synonymous or splicing mutant alleles by the regression model. OncodriveFML also detects in total 9 uniquely significant genes in all cancers, among which 7 (77.8%) genes are in the CGC gene list. Among the 9 genes, 5 genes have suggestive p-values (p<1E-4) by WITER. In the comparison, we ignore ITER because all significant genes by ITER are also significant by WITER.

### Rescued significant genes in smaller samples

We investigated the scenario in which the significant genes in a smaller sample missed by a tool can be rescued by another tool. We randomly drew six sub-samples of half size from the largest dataset, BRCA, and estimated cancer driver-genes by the three tools, MutSigCV, OncodriveFML and WITER. As shown in Table S8, using half samples, WITER misses only ∼22% genes on average which are significant in the full sample by the same method. The usage of MutSigCV and OncodriveFML in the sub-sample rescues almost none of the missed genes by WITER. MutSigCV misses ∼53% genes on average which are significant in the full sample by the same method. When the same half-size samples are analyzed by WITER, it rescues ∼42% of the missed genes by MutSigCV. The usage of OncodriveFML only rescues 6% of the missed genes by MutSigCV. OncodriveFML misses ∼61% genes on average which are significant in the full sample by the same method. When the same half-size samples are analyzed by WITER, it rescues ∼35% of the missed genes by OncodriveFML. The usage of MutSigCV rescues none of the missed genes by OncodriveFML. The higher proportions of significant genes and rescued genes by WITER in the sub-samples again shows that WITER has enhanced power to detect driver genes that would be missed by alternative methods.

### Performance in 23 cancer datasets with relatively small samples

Another important advantage of WITER is its ability to detect cancer-driver genes in small samples with a usage of reference samples. We applied the approach to other 23 cancers of small samples. We deliberately use two different reference cancers samples (BRCA and SKCM) with low and high background mutation rates to investigate whether WITER is sensitive to the reference samples. Three results are obtained. First, the usage of the reference datasets substantially improves the distribution of p-values. According to the QQ plots (Figure S1), the p-value distributions of the background genes (FWER>0.05) with reference samples are very close to the uniform distributions. In contrast, the p-values of the background genes without reference sample are weird and do not follow the uniform distribution. Second, WITER with reference samples detects significant genes even in cancers with very small sample size (See the results in Table 5). Among the 18 cancers with one or more significant genes (FWER=0.05), 3 cancers have less than 50 subjects, e.g., LB (n=26), LUSE (n=30), and CESC (n=37). Almost all significant genes are in CGC list. Third, it seems the difference in reference samples has a small and simple influence on the number of significant genes. The reference sample (BRCA) with low background mutation rate leads to slightly more significant genes than the one (SKCM) with high background mutation rate for the tested cancers. Moreover, we note that almost all significant genes according to the high background mutation rate reference sample are also significant according to the low background-mutation rate reference sample. Therefore, false positive findings can be easily controlled by using a high background mutation rate reference sample in practice although this may increase the false negatives. Anyhow, the overlapping of the significant genes according to the two extreme references is still high. In three cancers (LAML, LGG and GBM) with at least 5 significant genes, WITER using SKCM reference detects 90%, 100% and 100% overlapped significant genes with that using BRCA respectively.

**Table 5:** The number of significant genes in 23 cancers with two reference samples of different background mutations by WITER

|  | SKCM Ref |  | BRCA Ref |  | Overlapped Gene Number |
| --- | --- | --- | --- | --- | --- |
|  | Sig. Genes | CGC Genes | Sig. Genes | CGC. Genes |  |
| **LAML** | 10 | 10 | 13 | 12 | 9 |
| **LGG** | 7 | 7 | 7 | 7 | 7 |
| **GBM** | 6 | 6 | 7 | 7 | 6 |
| **MM** | 3 | 3 | 6 | 4 | 3 |
| **MED** | 4 | 4 | 4 | 4 | 3 |
| **PAAD** | 4 | 4 | 4 | 4 | 4 |
| **THCA** | 3 | 3 | 4 | 3 | 3 |
| **PRAD** | 3 | 3 | 3 | 3 | 3 |
| **CLL** | 2 | 2 | 3 | 3 | 2 |
| **DLBCL** | 2 | 1 | 3 | 2 | 2 |
| **LIHC** | 2 | 2 | 2 | 2 | 2 |
| **ESCA** | 1 | 1 | 2 | 2 | 1 |
| **CESC** | 0 | 0 | 2 | 0 | 0 |
| **KICH** | 1 | 1 | 1 | 1 | 1 |
| **LUSE** | 1 | 1 | 1 | 1 | 1 |
| **NB** | 1 | 1 | 1 | 1 | 1 |
| **LB** | 0 | 0 | 1 | 1 | 0 |
| **ALL** | 0 | 0 | 0 | 0 | 0 |
| **AT** | 0 | 0 | 0 | 0 | 0 |
| **CARC** | 0 | 0 | 0 | 0 | 0 |
| **KIRP** | 0 | 0 | 0 | 0 | 0 |
| **RHAB** | 0 | 0 | 0 | 0 | 0 |
| **STS** | 0 | 0 | 0 | 0 | 0 |

Note: Sig. Genes: Significant genes according to Bonferroni correction for family-wise error rate 0.05; CGC Genes: The significant genes in cancer gene consensus list; Overlapped Gene Number: the number of overlapped significant genes according to two different reference samples SKCM and BRCA. The full names of cancers are in Table S1

**Table 6:**
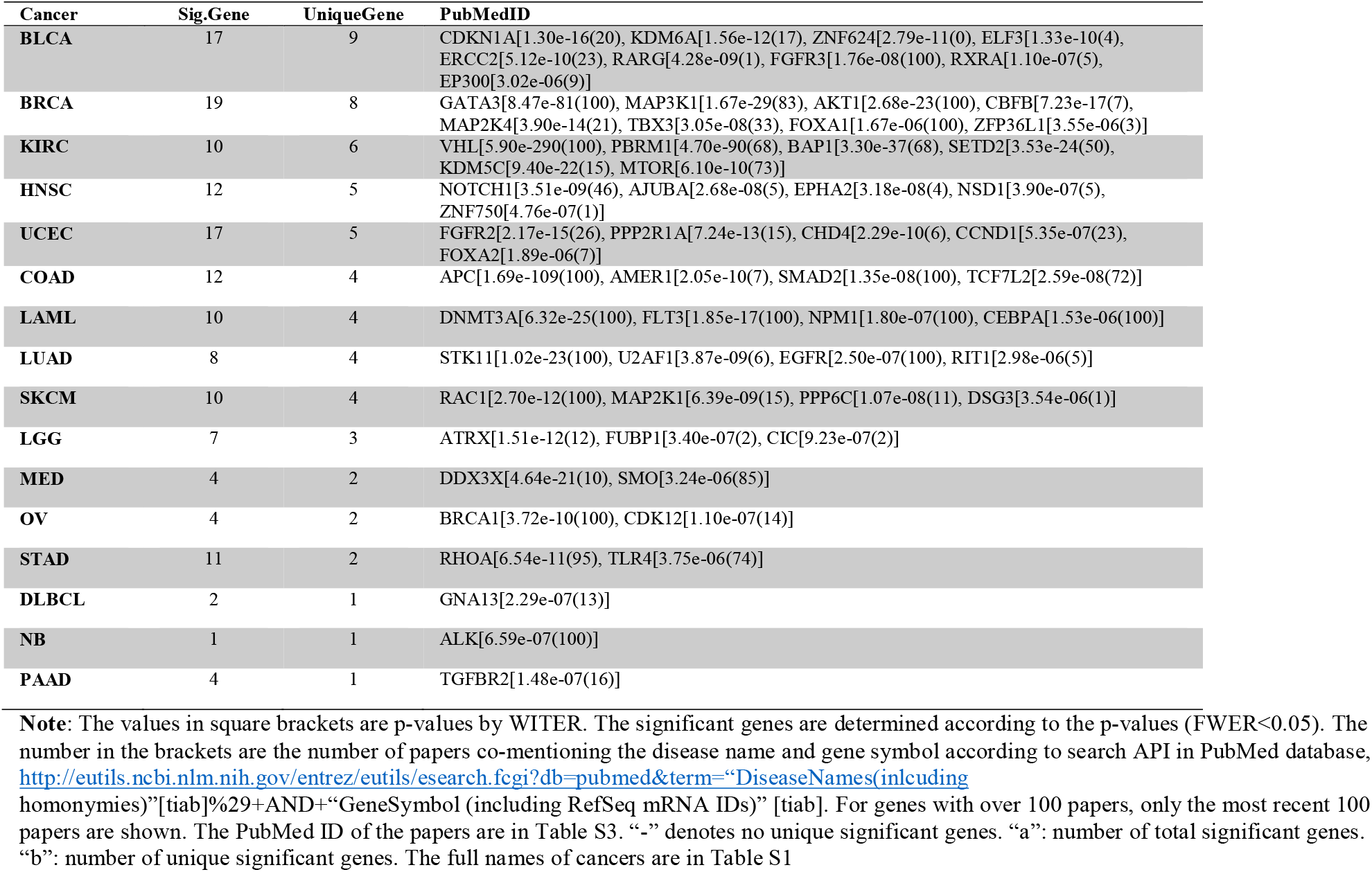
Genes significant only in one cancer and hit papers in PubMed database

**Table 6:** Predictors used in the regression model of estimating somatic mutations in passenger genes

| Predictor | Is used as cancer-specific | Sources | Note |
| --- | --- | --- | --- |
| **Length of genomic regions** | No | RefGene and GECODE | Combined coding regions |
| Gene expression in cancers tissues | Yes | https://tcga.xenahubs.net:443<br>TCGA Data Coordinating Center (DCC) | Batch effects normalized mRNA data called from RNAseq for each gene. |
| Gene expression in cancer cell lines | No | Curated by MutSigCV | Averaged expression across 91 cell lines in the Cancer Cell Line Encyclopedia |
| Chromatin accessibility by HiC data | No | Curated by MutSigCV | In K562 cells, ranging approximately from -50 (very closed) to +50 (very open) |
| Chromatin accessibility by ATAC-Seq data | Yes | https://atacseq.xenahubs.net | Quantile normalized log2[(ATAC-Seq counts+5)per million] |
| DNA-methylation level | Yes | https://atacseq.xenahubs.net | Beta values by DNA methylation HumanMethylation450 array |
| Gene's copy number variation in cancer tissues | Yes | https://atacseq.xenahubs.net | Gene-level copy number variation (CNV) estimated using the GISTIC2 threshold method. |
| DNA-replication timing | No | Curated by MutSigCV | HeLa cells, ranging from 100 (very early) to 1000 (very late) |
| Number of synonymous mutations | Yes | Local Samples | - |
| Constraint scores for non-synonymous mutations | No | Samocha et al (2014)(16) | Normalized student residues |

Note: The missing values of the predictors are imputed by a widely-used nonparametric method, missForest, in a R package.

It should be also noted that the additional significant genes according to the low background mutation rate reference sample are not necessarily spurious driver genes. For instance, WT1 is a significant driver gene of LAML based on the BRCA reference (p=3.33E-6) but insignificant based on SKCM reference(p=1.6E-4). WT1 is a well-known driver gene of acute lymphoblastic leukemia (24). To reduce possible false positive results rigorously, we use the results from the conservative reference sample, SKCM, for the subsequent analysis.

### Factors influencing power of WITER

We also investigated factors influencing the number of significant genes by WITER, which implies factors affecting its power. In a linear regression model, the sample size is strongly correlated with the number of significant genes, with a coefficient of determination R^2^, 0.34 (Figure S2). According to the fitted model, an estimation of sample size, ∼505, is required to detect 10 significant genes by WITER. Meanwhile, the number of significant genes is also related with the exome-wide background mutation rate, R^2^=0.17 (Figure S3). For example, the LUSC has a high genome-wide background mutation rate, 305 mutations per exome. A relatively smaller sample, 175, has led to detection of 8 significant genes. Therefore, when the sample size and number of mutations per sample enter as explanatory variables in a linear regression model, both have significant and positive correlation with the number of significant genes (p=1.77E-5 at the sample size and 0.0014 at number of mutations per exome). The coefficient of determination R^2^ increases to 0.54. After correcting for the mutation rate, the estimated sample size for 10 significant genes at LUSC changes from 505 to 199, which is closer to the real data. Table S1 lists the number of estimated sample size for detection of 10 significant genes by WITER at 11 cancers. While the three cancers types (BRCA, KIRC and OV) need a large sample size (more than 500 subjects) for detection of 10 significant driver genes by WITER, other three cancer types (LUSC, SKCM and LUAD) need only a sample size of less than 260.

### The landscape of driver-genes at 26 cancer types

WITER’s advantage of effectiveness in small samples enables the production of a comprehensive landscape of driver-genes in multiple cancer types. It detects one or more significant driver genes in 26 cancer types, out of which 13 cancer types have more than 5 genes (FWR<0.05, See details in Figure 4 and the Supplementary Excel File 1). The cancer with the largest sample, BRCA, has the largest number of significant genes, 19. Two cancers, UCEC and BLCA, which have relatively higher background mutation rates, have the second largest number of significant genes, 17. Twenty-seven genes occur in two or more cancers. However, the number of overlapped significant genes (Table S2) in the cancers are too small to produce sensible cancer clusters although we see cancers of similar origin have common genes (CLL and DLBC). As expected, the famous tumor suppressor gene, TP53, is the commonest significant genes (in 24 cancer types), followed by PIK3CA, PTEN, KRAS, NRAS and RB1, each of which is estimated as driver genes for 5 or more cancer types.

**Figure 4.**
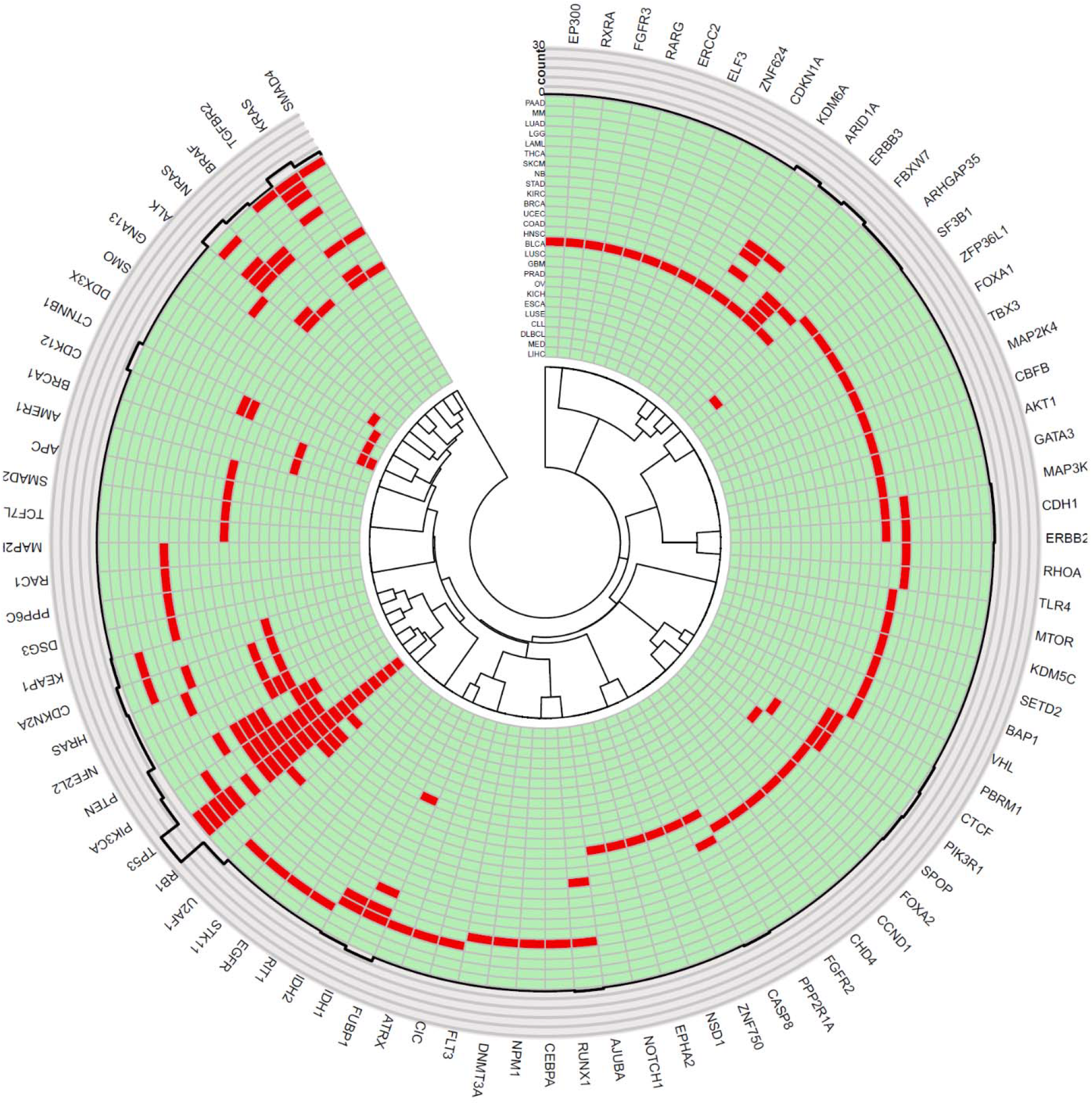
Circos plot displays 178 significant genes in 26 cancers. Notes: The innermost ring denotes dendrogram of genes. The next ring contains significant genes (marked in red) of corresponding cancers. It is followed by a ring of counts cancers in which the genes are significant. The outmost ring contains gene symbols. The full names of cancers are in Table S1.

Sixty-one genes are significant in only one out of the 26 cancers (See details in Table 4 and Table S5). Cancers with more significant genes tend to have more such uniquely significant genes, implying high genetic heterogeneity. In the *in-silico* validation analysis, 60 out of the 61 genes have one or more hit papers for the corresponding cancer types in the NCBI PubMed database (Table 6 and S3). Fifty-two (=85.2%) genes have more than 5 hit papers (Figure S6). There are even 17 genes having 100 or more hit papers. The high literature supporting rate suggests the high accuracy of WITER for estimating driver-genes of specific cancers even in small samples. It should be noted that the uniquely significant genes of a cancer do not necessarily mean uniquely driver genes of the cancer. As sample sizes increase, the genes may become significant in other cancer types. However, the uniquely significant genes of a cancer may imply relatively high contribution to the particular cancer when the sample size is not large. For instance, the BRCA has 8 uniquely significant genes. Six out of the 8 genes have at least 20 hit papers in the NCBI PubMed database, an indication of the strong relationship between the genes and BRCA. Interestingly, although BRCA1 is an important tumor suppressor gene for familiar BRCA, it is uniquely significant in OV. While BRCA1 has an insignificant p-value (p=1.38E-4) in the BRCA dataset, it achieves a p-value 3.72E-10 in the OV dataset. BRCA1 also has 100 hit papers for OV in the NCBI PubMed database. Probably, for sporadic cases, BRCA1 is a stronger driver for OV than BRCA.

There are 7 uniquely significant genes having only three or fewer hit papers in NCBI PubMed database. The BLCA has two such genes, RARG and ZNF624. RARG encodes a retinoic acid receptor that belongs to the nuclear hormone receptor family. This gene has been linked to leukemia repeatedly by multiple studies (e.g. (25,26)). This is the first time that it is suggested as a driver gene of BLCA. ZNF624 encodes a zinc finger protein 624. It has 19 non-synonymous mutations and a high functional gene score 42 (p= 2.79E-11 by WITER). It seems that this gene has not been well-studied yet. There are few papers mentioning this gene in the PubMed database. The uniquely significant gene of BRCA, ZFP36L1, has three hit papers. ZFP36L1 encodes a putative nuclear transcription factor that likely functions in regulating the response to growth factors. A recent study suggests that inactivating phosphorylation of ZFP36L1 is related to promoted specification of the triple-negative breast cancer phenotype, a subtype cancer of poor prognosis (27). A uniquely significant gene of HNSC, ZNF750, only has one hit paper. ZNF750 also encodes a zinc finger protein. The hit paper reported a down regulation of ZNF750 by gene fusions in human papillomavirus (HPV) positive HNSC samples(28). ZNF750 has been suggested to be associated with esophageal squamous cell carcinoma (29), which indirectly supports its cancer driver potential to HNSC. The uniquely significant gene of SKCM, DSG3, only has one hit paper. DSG3 encodes a member of the desmoglein family and cadherin cell adhesion molecule superfamily of proteins. Although there have been no results directly linking DSG3 to SKCM, multiple studies suggest that DSG3 may activate Src signaling, an important pathway in cancer invasion, e.g., (30). The last two genes, FUBP1 and CIC, are uniquely significant for LGG. FUBP1 encodes a single stranded DNA-binding protein that binds to multiple DNA elements, including the far upstream element (FUSE) located in upstream of c-myc. FUBP1, has two hit papers on LGG in the PubMed database. One of the papers reported evidences of FUBP1’s role in neuronal differentiation and explained its tumor-suppressor function in the nervous system(31). CIC encodes capicua transcriptional repressor and has two hit papers in the PubMed database for LGG. One of papers suggests that CIC may work with ATXN1-ATXN1L as a potent regulator of the cell cycle related with development of cancers(32). These literature evidences strongly support these most of the 7 genes are potential driver genes (although relatively new) for the corresponding cancers.

Among the above 7 uniquely significant genes, five (ZNF624, RARG, ZFP36L1, ZNF750 and DSG3) also have estimation results by MutSigCV and OncodriveFML. It turns out all the 5 genes have insignificant p-values by MutSigCV and OncodriveFML in the corresponding cancers. For the remaining two genes, FUBP1 and CIC, because the number of total variant number is too small, we do not perform the driver gene estimation analysis by MutSigCV and OncodriveFML. WITER has a unique advantage of being workable for small sample and detects the two potential driver genes for LGG.

## Discussion

Accurately modeling counts of somatic mutations at background genes has long been a fundamental technical challenge in genomic characterization of cancer-driver genes (2,11). The proposed approach, WITER, has four technical advances to address this issue. The first one is the advanced model, ZTNB regression, that fits the number of somatic mutations at background genes better. In samples of a typic size, one often sees an inflation of zero mutation genes and overdispersion of mutation counts. The inflated zero values make it difficult to fit the distribution of genomic counts based on conventional distributions. The ZTNB distribution adequately addresses the zero inflation and the overdispersion issues. This is demonstrated by the results that ZTNB model always achieves the minimal AIC among four alternative models (Table 2) and statistically valid *p* values distribution(Figure 2 and S1). The valid *p* values distribution solves the common problem in alternative methods that resort to time-consuming simulation or permutation for statistical inference. The other three technical advances include iterative regression, integrating reference samples and imposing prior weights at variants. The iterative regression relieves distortion of driver genes to the background baseline so that the residues of mutation counts at driver genes are not shrunk. Its allowance of integrating reference samples ensures a stable resulting model and thus a valid estimation in small sample. We find that the number of significant genes detected by WITER is generally not sensitive to the reference samples in most cancers (Table 5). We show that imposing prior weights at variants enables detection of more cancer consensus driver genes in all the tested datasets. These four technical advances together determine the enhanced power of WITER for detecting more cancer-driver genes than alternative methods while effectively controlling statistical type 1 error.

The ZTNB model and three-tier framework are generic and can be extended to other types of mutations and different predictors. In the present paper, we focus on the non-synonymous and splicing variants (including SNVs and InDels). This is because the availability of abundant data (e.g., exome sequencing data) in the public domains greatly facilitates the methodological validation. It is true that non-coding variants (usually discovered by whole genome sequencing) also contribute to development of cancers(33). However, the public resources of whole-genome sequencing are much fewer than that of the exome sequencing. Theoretically, one can replace the non-synonymous variants with non-coding variants like upstream-or-downstream variants while the predictors are replaced correspondingly. As long as the counts of non-coding variants have three characteristics, zero-inflation, over-dispersion, majority of null hypotheses, the key principle and main frame should also be workable. As the cost of high-coverage whole genome sequencing is decreasing, more data will be available for an evaluation of the method in non-coding variants in the future.

A slightly unexpecting finding is that some genomic features (e.g., gene expression and chromatin accessibility) from cancer non-specific cell lines are generally more relevant to somatic mutations than that from cancer-matched primary tumors. Lawrence et al. (2013) stated that matched normal tissues led to similar results as the cancer non-specific cell lines for the correlation between somatic mutation frequency in cancers and gene expression level (6). The present study uses matched cancer tissues instead of matched normal tissues, which may be the cause of the difference. Polak et al (2015) suggested cell-of-origin chromatin features (including chromatin accessibility) are stronger determinants of cancer mutation profiles of the entire genome than chromatin features of matched cancer cell lines (12). In the present study, our finding exclusively used chromatin accessibility in coding regions, which is not equivalent to Polak et al (2015). Although the underlying causes of the differences are subject to more and deeper study in the future, the strong correlation between somatic mutations and the cancer non-specific predictors makes WITER more flexible in practice.

We compared the proposed method with two widely-used and well-performed approaches(11), both of which belong to the unsupervised category. Another category of methods is the supervised approaches for detecting cancer driver genes. According to Tokheim et al (2016)(11), the supervised method 20/20plus outperformed the best unsupervised methods at that time (including MutSigCV and OncodriveFML) in terms of p-value distributions and the number of significant genes. However, a supervised strategy has learning bias toward the training samples in nature(34). If the training sample is not representative of all sample, the trained model will have low power in new samples. This would be particularly true for cancers because of their high genetic heterogeneity(5). Second, the 20/20plus also used many common genomic features of a gene (e.g., evolutionary conservation, predicted functional impact of variants, and gene interaction network connectivity) in the prediction(11). Although the usage of common genomic features will add information to prioritize common cancer-driver genes, it also runs the risk of diluting the information in local sample for identifying unique cancer driver genes, which would be important for a precision diagnosis and treatment of a specific cancer. Finally, the 20/20plus resorted time-consuming permutation procedure to generate p-values for statistical test. In contrast, the WITER and ITER are much faster than 20/20plus because it calculates p-values in an analytic way. Nevertheless, we also made additional comparisons between WITER and 20/20plus approach in the 11 cancers. The p-value distributions of background ground genes produced by both methods are similar and approximately follow uniform distribution. (See QQ plots in Figure S4). While 20/20plus detected more significant and cancer-consensus genes in 5 out of the 11 cancers (See details in Figure S5), WITER detected more significant and cancer-consensus genes in other 5 cancers. They rescued similar number of missed genes for each other in an experiment(See details in Table S8).

A limitation of the present study is that many true cancer-driver genes are generally unknown for most cancers. Most results inflecting enhanced power of WITER in the paper are indirect. We did not simulate datasets to quantify the power difference of WITER and alternative methods by artificially setting “true” driver genes because there are many unknown factors shaping the landscape of somatic mutations. An artificial model in simulation is often too subjective to reflect the reality. The usage of real data and *in-silico* validation are widely adopted and often effective in methodological studies(11). In the present study we showed WITER detects more significant genes than alternative methods in all 11 tested cancers. A high coverage of these genes in CGC list and co-mentioned with the corresponding cancers in titles or abstracts of more than 5 papers in PubMed database is shown. The enhanced power of WITER is further confirmed by more rescued genes for alternative methods in half-size samples. These results strongly suggest that there is a high true discovery rate in the significant genes by WITER and it detects more genuine cancer-driver genes than alternative methods.

Applying the powerful approach, WITER, we generated a landscape of driver genes in 26 cancers. Its unique ability of integrating reference sample enables detection of driver genes in samples of size as small as 30 although more driver genes will be detected in larger samples. The analysis revealed many genes which are common driver genes for multiple cancers. Most majority of the genes have many literature supports. The common driver genes may be effective drug targets for treatment of the cancers. Meanwhile, there are also a lot of significant genes which are unique for a single cancer. Some of these genes may be unique driver genes of the corresponding cancers although sample sizes of other cancers may change the results. The unique driver genes are potentially effective for a precision diagnosis and treatment of corresponding tumors. In the results section, we highlight 7 uniquely significant genes by WITER in 5 different cancers. These genes have three or fewer hit papers in the PubMed database which may be potential new driver genes for corresponding cancers. In a detailed literature survey, these genes have very strong implication to development of cancers. In contrast, all the 7 genes are either insignificant by MutSigCV and OncodriveFML or cannot be tested due to small sample size.

## Acknowledgements

This work was funded by National Natural Science Foundation of China (31771401), Science and Technology Program of Guangzhou (201803010116), Hong Kong Health and Medical Research Fund (02132236). Hong Kong General Research Fund 17124017, 17121414 and TRS T12C-714/14-R. We thank Tokheima and colleagues for sharing the high-quality curated somatic mutations in 32 cancers from multiple resources.

