## Supplementary Notes and Figures and Tables for "WITER: A powerful method for the estimation of cancer-driver genes using a weighted iterative regression accurately modelling background mutation rate"

**Calculation of deviance residue of ZTNB regression**

The coefficients can be estimated by maximum likelihood with a quasi-Newton method. In our study, we called the maximum likelihood method in a R package countreg (<https://r-forge.r-project.org/R/?group_id=522>) to estimate the coefficients. The dispersion parameter $\theta$ is jointly estimated with the regression coefficients, $\beta_{0},\cdots,\beta_{6}$. The model is fitted only for genes with non-zero counts.

With the established model, the logarithm of the expected mutation counts, $log{(\hat{\mu}}_{i})$, at somatic variants in a gene *i* can be calculated by:

$$log{(\hat{\mu}}_{i})=\hat{\beta}_{0}+\hat{\beta}_{1}x_{i,1}+\hat{\beta}_{2}x_{i,2}+\hat{\beta}_{3}x_{i,3}+\hat{\beta}_{4}x_{i,4}+\hat{\beta}_{5}x_{i,5}+\hat{\beta}_{6}x_{i,6}$$

where $\hat{\beta}_{0},\cdots,\hat{\beta}_{6}$ are the estimated coefficients.

Given the fitted parameters, the probability of zero mutation gene *i* is: $p_{i, 0}=\left( \frac{\hat{\theta}}{\hat{\theta}+\hat{\mu}_{i}} \right)^{\hat{\theta}}$.

Under zero-truncated model, the raw residual at gene *i* is:

$r_{i}=y_{i}-\frac{\hat{\mu}_{i}}{1-p_{i, 0}}$.

The deviance residual of the model at gene *i* is:

$e_{i}=sign(r_{i})*\sqrt{2*|ll({y_{i}|\mu}_{i}^{*},\hat{\theta})-ll(y_{i}|\hat{\mu}_{i},\hat{\theta})|}$ ,

where $sign(x)$ is the standard sign function, $ll(\mu,\theta)$ is the natural logarithm of the likelihood function of the zero-truncated negative binomial distribution,

$ll(y_{i}|\mu,\theta)=ln[g(y_{i}|\mu,\theta)]$,

and $\mu_{i}^{*}$ is the estimated mean given the observed count $y_{i}$ and estimated $\hat{\theta}$ of a saturated model, obtained by solving the following equation:

$$y_{i}=\frac{\mu_{i}^{*}}{1-\left( \frac{\hat{\theta}}{\hat{\theta}+\mu_{i}^{*}} \right)^{\hat{\theta}}}$$

The deviance residuals are further standardized by the estimated mean $\hat{\mu}_{e}$ and standard deviation $\hat{\sigma}_{e}$ of the deviance residuals,

$\acute{e}_{i}=\frac{e_{i}-\hat{\mu}_{e}}{\hat{\sigma}_{e}}$.

**Construction of prior weights at variants**

A random forest model was trained by COSMIC database to generate prior weights at variants. To avoid circular bias, all subjects (n=7,916) in our collected testing samples of the 34 cancers were excluded from COSMIC database. In COSMIC(V83), we collected 27,09 somatic mutation variants occurring over 20 times in primary cancer tissues to constitute a positive variant set. A negative control variant set consists of 863,304 somatic mutation variants which occurred only once in primary cancer tissues. Because the negative set is much larger than the positive set, the final prediction model is an ensemble of 10 random forest models by down-sampling in the negative dataset to match the equal sample sizes in positive and negative datasets. The predictors at each variant include 19 deleterious or conservation scores from the database dbNSFP v3.5 (13), (e.g., MutationTaster2 (14) and FATHMM (15), see the names of all tools in Supplementary Figure S7). The area under the receiver operating characteristic curve of the random forest model is 83%, which is better than a multivariate logistic regression model and individual predictors (Figure S7). The random forest prediction scores, *s*, ranged from 0 to 1. For variants without prediction scores due to missing values, the average score in the gene is used.

**The usage of the alternative driver-gene estimation methods**

MutSigCV is a powerful method for detecting genes mutated more often than expected by chance. It used a local regression model to estimate the expected mutant alleles by multiple genomic features of a gene in cancer cells including its expression level, replication time and 3D chromatin interaction capture (HiC). The online MutsigCV version (1.2) was used through the Broad website (<http://genepattern.broadinstitute.org/gp/pages/index.jsf?lsid=MutSigCV>). The recommended exome coverage file (<https://genepattern.broadinstitute.org/gp/data/xchip/gpprod/shared_data/example_files/MutSigCV_1.3/exome_full192.coverage.txt>) and gene covariates file ([https://genepattern.broadinstitute.org/gp/data//xchip/gpprod/shared_data/example_files/MutSigCV_1.3/gene.covariates.txt](https://genepattern.broadinstitute.org/gp/data/xchip/gpprod/shared_data/example_files/MutSigCV_1.3/gene.covariates.txt)) were used. OncodriveFML is a method designed to estimate the accumulated functional impact bias of tumor somatic mutations in both coding and non-coding genomic regions, based on a simulation process. It used CADD scores to predict mutational impacts. The results were produced according to coding DNA sequence (CDS) regions. The genome reference and CDS files were downloaded from the website (<https://bitbucket.org/bbglab/oncodrivefml>) as the authors recommended. The default parameters of OncodriveFML were used to produce the results. The 20/20 plus is a machine-learning-based method integrating multiple features to predict driver genes, including sample mutational clustering, evolutionary conservation, predicted functional impact of variants, mutation consequence types, gene interaction network connectivity, etc. It used computer simulation to generate p-values for statistical significance. The 20/20plus v1.1.3 was downloaded and installed according to the website tutorial (<http://2020plus.readthedocs.io/en/latest/index.html>). The necessary files were also collected as the authors suggested (<http://probabilistic2020.readthedocs.io/en/latest/tutorial.html#gene-bed-file> and <http://probabilistic2020.readthedocs.io/en/latest/tutorial.html#pre-computed-scores-optional>). The data were analyzed by a pipeline to predict the cancer drivers under the default parameters. The 20/20 plus took 1.5 hours on average to analyze a dataset on a computer with 12 CPU (1.70GHz) cores and 64G RAM. The number of simulations was 10000.

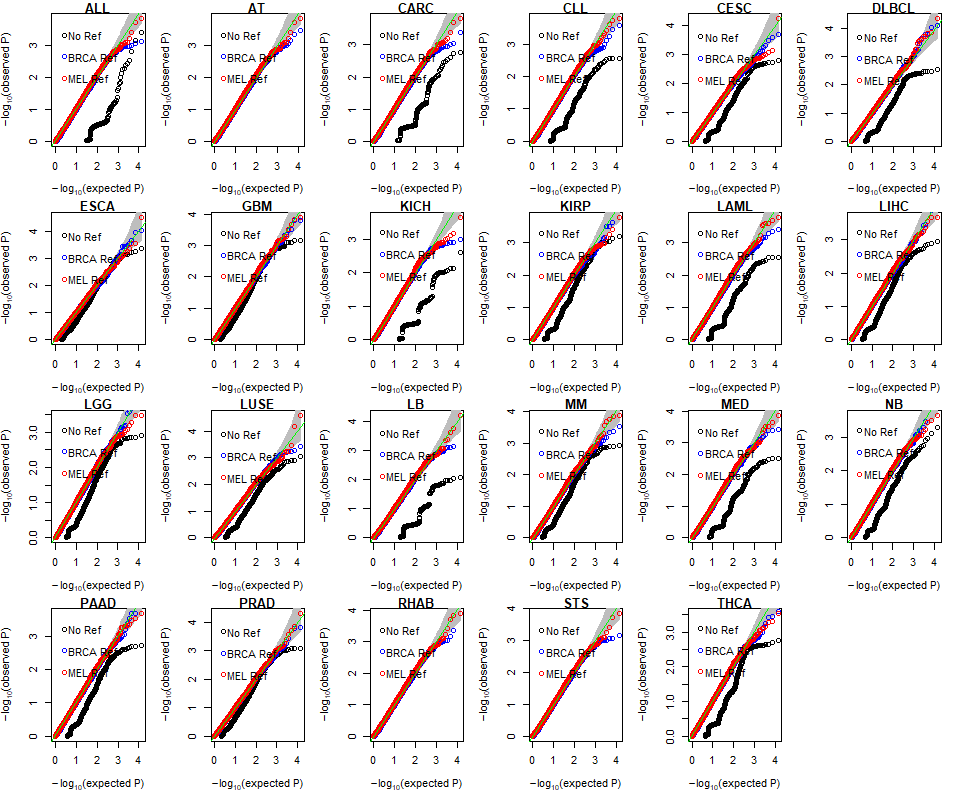

Figure S1: the QQ plots of p-values in 23 sample by WITER with and without reference samples

Note: The p-values less than a cutoff according to FWER 0.05 were excluded.

|  | SampleSize |
| --- | --- |
| 394 | 8 |
| 289 | 10 |
| 407 | 12 |
| 1093 | 19 |
| 175 | 8 |
| 244 | 11 |
| 255 | 17 |
| 142 | 17 |
| 244 | 12 |
| 484 | 10 |
| 480 | 4 |
| 365 | 6 |
| 160 | 1 |
| 420 | 3 |
| 205 | 3 |
| 227 | 7 |
| 30 | 1 |
| 233 | 4 |
| 150 | 2 |
| 111 | 0 |
| 37 | 0 |
| 325 | 3 |
| 56 | 2 |
| 351 | 1 |
| 196 | 10 |
| 331 | 4 |
| 223 | 2 |
| 54 | 0 |
| 65 | 1 |
| 26 | 0 |
| 55 | 0 |
| 32 | 0 |
| 15 | 0 |
| 42 | 0 |

**Figure S2:** The linear regression between number of significant genes and sample size
The dashed line is the fitted line. R2 is the coefficient of determination.

|  | Number of somatic mutation variants (Thousand) |
| --- | --- |
| 269.259 | 8 |
| 327.734 | 10 |
| 145.919 | 12 |
| 50.258 | 19 |
| 305.051 | 8 |
| 172.594 | 11 |
| 151.894 | 17 |
| 236.782 | 17 |
| 128.939 | 12 |
| 58.2624 | 10 |
| 58.2208 | 4 |
| 59.1808 | 6 |
| 118.344 | 1 |
| 39.5667 | 3 |
| 52.0146 | 3 |
| 42.3789 | 7 |
| 279.233 | 1 |
| 32.9356 | 4 |
| 50.76 | 2 |
| 67.6847 | 0 |
| 164.973 | 0 |
| 18.7538 | 3 |
| 102.536 | 2 |
| 14.3647 | 1 |
| 20.6735 | 10 |
| 10.5227 | 4 |
| 15.4933 | 2 |
| 30.5556 | 0 |
| 19.4308 | 1 |
| 44.9231 | 0 |
| 11.1455 | 0 |
| 7.5 | 0 |
| 7.8 | 0 |
| 2.2619 | 0 |

**Figure S3:** The linear regression between number of significant genes and number of somatic mutation variants
The dashed line is the fitted line. R2 is the coefficient of determination.

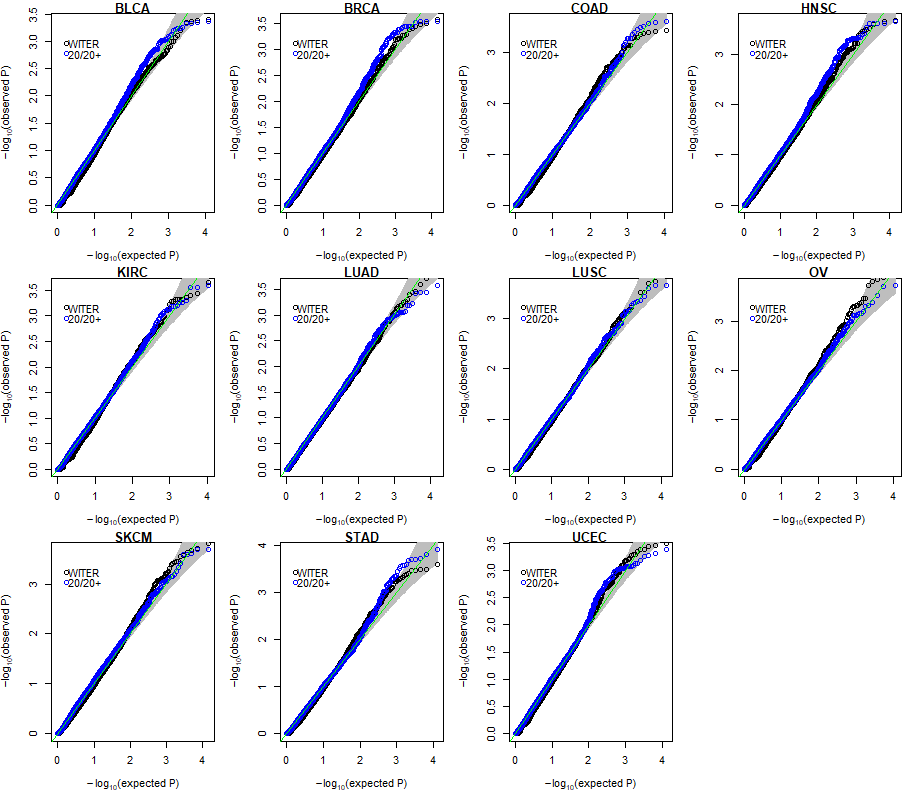

Figure S4. QQ plot of background gene p-values produced by WITER and 2020 plus methods in 11 cancers. The p-values less than a cutoff according to FWER 0.05 were excluded.

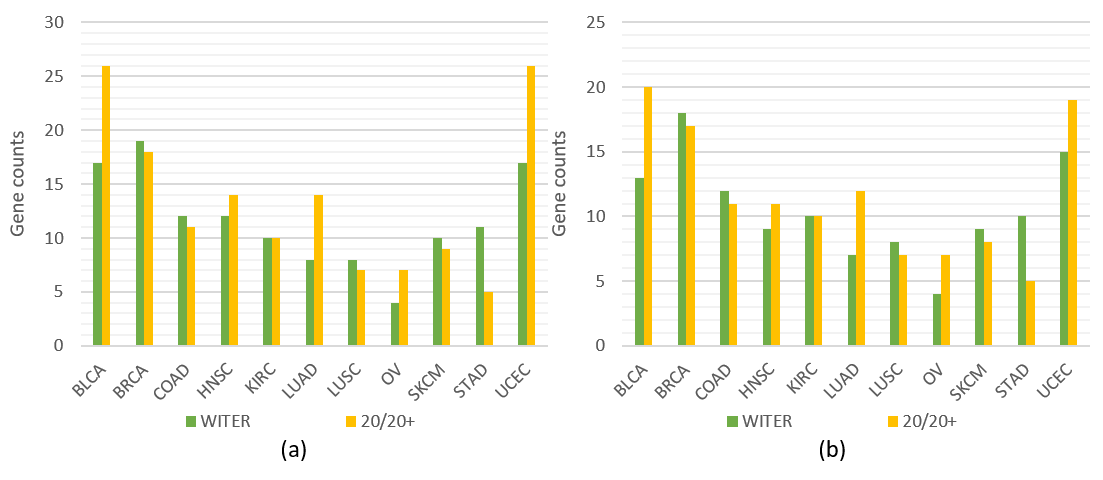

**Figure S5:** Performance comparison of WITER and 20/20+ for detecting cancer driver mutation in 11 cancers
a: the number of significant genes; b: cancer consensus significant genes. The p-values less than a cutoff according to FWER 0.05 were excluded.

Accumulated proportion

| PubMed hits | Unique |
| --- | --- |
| 0 | 1 |
| 5 | 0.852459 |
| 10 | 0.688525 |
| 15 | 0.606557 |
| 20 | 0.52459 |
| 25 | 0.459016 |
| 30 | 0.442623 |
| 35 | 0.42623 |
| 40 | 0.42623 |
| 45 | 0.42623 |
| 50 | 0.409836 |
| 55 | 0.393443 |
| 60 | 0.393443 |
| 65 | 0.393443 |
| 70 | 0.360656 |
| 75 | 0.311475 |
| 80 | 0.311475 |
| 85 | 0.295082 |
| 90 | 0.278689 |
| 95 | 0.278689 |

Figure S6: The accumulated proportion of uniquely significant genes with PubMed hit papers.

area under curve of the receiver operating characteristic (ROC AUC) curve

| Prediction model | 系列 1 |
| --- | --- |
| SIFT | 0.525027 |
| Polyphen2_HDIV | 0.522917 |
| Polyphen2_HVAR | 0.510589 |
| LRT | 0.510299 |
| MutationTaster | 0.493384 |
| MutationAssessor | 0.505971 |
| FATHMM | 0.512379 |
| PROVEAN | 0.51072 |
| VEST3 | 0.516222 |
| M-CAP | 0.498458 |
| MutPred | 0.504027 |
| CADD_raw | 0.503573 |
| DANN | 0.506273 |
| fathmm-MKL_coding | 0.505802 |
| Eigen-raw | 0.526664 |
| GenoCanyon | 0.526588 |
| integrated_fitCons | 0.513634 |
| GERP++_RS | 0.530552 |
| SiPhy_29way_logOdds | 0.523559 |
| LogisticRegression | 0.781543 |
| RandomForest | 0.827208 |

Figure S7: Performance of predicting frequent mutant variants by individual and combined methods. The 5 folder-cross validation was used to generate the AUC. The evaluation was carried out by a Java package WEKA(V3, https://www.cs.waikato.ac.nz/ml/weka/)

Table S1: Sample size and variant number of 34 cancer datasets

| Full Name | Abbreviation | Sample Size | Variant Number | Ratio | ^a^Estimated Sample Size | Search terms in PubMed Database |
| --- | --- | --- | --- | --- | --- | --- |
| Lung adenocarcinoma | LUAD | 394 | 106088 | 269.26 | 253 | Lung Adenocarcinoma;Lung Cancer |
| Skin Cutaneous Melanoma | SKCM | 289 | 94715 | 327.73 | 164 | Melanoma;Skin Cutaneous Melanoma |
| Head and Neck squamous cell carcinoma | HNSC | 407 | 59389 | 145.92 | 441 | Head and Neck Squamous Cell Carcinoma |
| Breast invasive carcinoma | BRCA | 1093 | 54932 | 50.26 | 587 | Breast Cancer;Breast Adenocarcinoma |
| Lung squamous cell carcinoma | LUSC | 175 | 53384 | 305.05 | 199 | Lung Squamous Cell Carcinoma;squamous cell carcinoma of the lung |
| Stomach adenocarcinoma | STAD | 244 | 42113 | 172.59 | 400 | Stomach Adenocarcinoma;Stomach cancer;gastric cancer |
| Uterine Corpus Endometrial Carcinoma | UCEC | 255 | 38733 | 151.89 | 432 | Endometrial Carcinoma;Endometrial Cancer;Uterine Corpus Endometrial Carcinoma |
| Bladder Urothelial Carcinoma | BLCA | 142 | 33623 | 236.78 | 303 | Bladder Urothelial Carcinoma;Urothelial Carcinoma;Bladder cancer |
| Colon adenocarcinoma | COAD | 244 | 31461 | 128.94 | 467 | Colon adenocarcinoma;colon cancer;colorectal cancer |
| Kidney renal clear cell carcinoma | KIRC | 484 | 28199 | 58.26 | 574 | Kidney Clear Cell Carcinoma;clear cell renal cell carcinoma |
| Ovarian serous cystadenocarcinoma | OV | 480 | 27946 | 58.22 | 574 | Ovarian serous cystadenocarcinoma;Ovarian Cancer;serous cystadenoma |
| Glioblastoma multiforme | GBM | 365 | 21601 | 59.18 | - | Glioblastoma Multiforme |
| Esophageal carcinoma | ESCA | 160 | 18935 | 118.34 | - | Esophageal Adenocarcinoma;Esophageal Cancer |
| Prostate adenocarcinoma | PRAD | 420 | 16618 | 39.57 | - | Prostate Adenocarcinoma;Prostate Cancer |
| Multiple myeloma | MM | 205 | 10663 | 52.01 | - | Multiple myeloma;plasma cell myeloma |
| Brain Lower Grade Glioma | LGG | 227 | 9620 | 42.38 | - | Low Grade Glioma;Low-grade glioma |
| Small cell lung carcinoma | LUSE | 30 | 8377 | 279.23 | - | Lung Small Cell Carcinoma;Small cell lung cancer |
| Pancreatic adenocarcinoma | PAAD | 233 | 7674 | 32.94 | - | Pancreatic Adenocarcinoma;Pancreatic Cancer |
| Liver hepatocellular carcinoma | LIHC | 150 | 7614 | 50.76 | - | Liver Hepatocellular carcinoma;Hepatocellular Carcinoma;Liver Cancer |
| Kidney renal papillary cell carcinoma | KIRP | 111 | 7513 | 67.68 | - | Kidney Papillary Cell Carcinoma;Papillary renal cell carcinoma |
| Cervical squamous cell carcinoma and endocervical adenocarcinoma | CESC | 37 | 6104 | 164.97 | - | Cervical Carcinoma |
| Thyroid carcinoma | THCA | 325 | 6095 | 18.75 | - | Thyroid Carcinoma;Thyroid Cancer |
| Diffuse large B-cell lymphoma | DLBCL | 56 | 5742 | 102.54 | - | Diffuse large B-cell lymphoma |
| Neuroblastoma | NB | 351 | 5042 | 14.36 | - | Neuroblastoma |
| Acute Myeloid Leukemia | LAML | 196 | 4052 | 20.67 | - | Acute myeloid leukemia |
| Medulloblastoma | MED | 331 | 3483 | 10.52 | - | Medulloblastoma |
| Chronic lymphocytic leukemia | CLL | 223 | 3455 | 15.49 | - | Chronic lymphocytic leukemia |
| Carcinoid Cancer | CARC | 54 | 1650 | 30.56 | - | Carcinoid Cancer |
| Kidney Chromophobe | KICH | 65 | 1263 | 19.43 | - | Kidney Chromophobe;renal chromophobe cell carcinoma |
| B-cell lymphomas | LB | 26 | 1168 | 44.92 | - | Lymphoma B-cell;B-cell lymphomas |
| Acute lymphoblastic leukemia | ALL | 55 | 613 | 11.15 | - | Acute lymphocytic leukemia |
| Rhabdoid tumor | RHAB | 32 | 240 | 7.5 | - | Rhabdoid tumor |
| Soft Tissue Sarcoma | STS | 15 | 117 | 7.8 | - | Soft Tissue Sarcoma;soft-tissue sarcoma;sarcoma |
| Astrocytoma | AT | 42 | 95 | 2.26 | - | Astrocytoma |

Note: The cancers in red are the 11 cancers with more total variants and used for the performance comparison. a: the estimated sample size is for detecting 30 significant genes by WITER. –: the sample sizes are not estimated because of the unreliable ratio derived in small samples.

Table S2: The overlapped significant genes among multiple cancers detected by WITER

|  | ALL | AT | BLCA | BRCA | CARC | CLL | CESC | COAD | DLBCL | ESCA | GBM | HNSC | KIRC | KICH | KIRP | LAML | LUAD | LUSC | LIHC | LGG | LUSE | LB | MM | MED | NB | OV | PAAD | PRAD | RHAB | SKCM | STAD | STS | THCA | UCEC |
| --- | --- | --- | --- | --- | --- | --- | --- | --- | --- | --- | --- | --- | --- | --- | --- | --- | --- | --- | --- | --- | --- | --- | --- | --- | --- | --- | --- | --- | --- | --- | --- | --- | --- | --- |
| ALL | 0 | - | - | - | - | - | - | - | - | - | - | - | - | - | - | - | - | - | - | - | - | - | - | - | - | - | - | - | - | - | - | - | - | - |
| AT | 0 | 0 | - | - | - | - | - | - | - | - | - | - | - | - | - | - | - | - | - | - | - | - | - | - | - | - | - | - | - | - | - | - | - | - |
| BLCA | 0 | 0 | 17 | - | - | - | - | - | - | - | - | - | - | - | - | - | - | - | - | - | - | - | - | - | - | - | - | - | - | - | - | - | - | - |
| BRCA | 0 | 0 | 3 | 19 | - | - | - | - | - | - | - | - | - | - | - | - | - | - | - | - | - | - | - | - | - | - | - | - | - | - | - | - | - | - |
| CARC | 0 | 0 | 0 | 0 | 0 | - | - | - | - | - | - | - | - | - | - | - | - | - | - | - | - | - | - | - | - | - | - | - | - | - | - | - | - | - |
| CLL | 0 | 0 | 1 | 2 | 0 | 2 | - | - | - | - | - | - | - | - | - | - | - | - | - | - | - | - | - | - | - | - | - | - | - | - | - | - | - | - |
| CESC | 0 | 0 | 0 | 0 | 0 | 0 | 0 | - | - | - | - | - | - | - | - | - | - | - | - | - | - | - | - | - | - | - | - | - | - | - | - | - | - | - |
| COAD | 0 | 0 | 3 | 2 | 0 | 1 | 0 | 12 | - | - | - | - | - | - | - | - | - | - | - | - | - | - | - | - | - | - | - | - | - | - | - | - | - | - |
| DLBCL | 0 | 0 | 1 | 1 | 0 | 1 | 0 | 1 | 2 | - | - | - | - | - | - | - | - | - | - | - | - | - | - | - | - | - | - | - | - | - | - | - | - | - |
| ESCA | 0 | 0 | 1 | 1 | 0 | 1 | 0 | 1 | 1 | 1 | - | - | - | - | - | - | - | - | - | - | - | - | - | - | - | - | - | - | - | - | - | - | - | - |
| GBM | 0 | 0 | 3 | 5 | 0 | 1 | 0 | 2 | 1 | 1 | 6 | - | - | - | - | - | - | - | - | - | - | - | - | - | - | - | - | - | - | - | - | - | - | - |
| HNSC | 0 | 0 | 4 | 3 | 0 | 1 | 0 | 3 | 1 | 1 | 2 | 12 | - | - | - | - | - | - | - | - | - | - | - | - | - | - | - | - | - | - | - | - | - | - |
| KIRC | 0 | 0 | 3 | 3 | 0 | 1 | 0 | 2 | 1 | 1 | 3 | 2 | 10 | - | - | - | - | - | - | - | - | - | - | - | - | - | - | - | - | - | - | - | - | - |
| KICH | 0 | 0 | 1 | 1 | 0 | 1 | 0 | 1 | 1 | 1 | 1 | 1 | 1 | 1 | - | - | - | - | - | - | - | - | - | - | - | - | - | - | - | - | - | - | - | - |
| KIRP | 0 | 0 | 0 | 0 | 0 | 0 | 0 | 0 | 0 | 0 | 0 | 0 | 0 | 0 | 0 | - | - | - | - | - | - | - | - | - | - | - | - | - | - | - | - | - | - | - |
| LAML | 0 | 0 | 1 | 2 | 0 | 1 | 0 | 3 | 1 | 1 | 2 | 1 | 1 | 1 | 0 | 10 | - | - | - | - | - | - | - | - | - | - | - | - | - | - | - | - | - | - |
| LUAD | 0 | 0 | 1 | 1 | 0 | 1 | 0 | 2 | 1 | 1 | 1 | 2 | 1 | 1 | 0 | 2 | 8 | - | - | - | - | - | - | - | - | - | - | - | - | - | - | - | - | - |
| LUSC | 0 | 0 | 4 | 4 | 0 | 1 | 0 | 2 | 1 | 1 | 4 | 5 | 3 | 1 | 0 | 1 | 3 | 8 | - | - | - | - | - | - | - | - | - | - | - | - | - | - | - | - |
| LIHC | 0 | 0 | 1 | 1 | 0 | 1 | 0 | 2 | 1 | 1 | 1 | 1 | 1 | 1 | 0 | 1 | 1 | 1 | 2 | - | - | - | - | - | - | - | - | - | - | - | - | - | - | - |
| LGG | 0 | 0 | 2 | 2 | 0 | 1 | 0 | 2 | 1 | 1 | 3 | 2 | 2 | 1 | 0 | 3 | 1 | 2 | 1 | 7 | - | - | - | - | - | - | - | - | - | - | - | - | - | - |
| LUSE | 0 | 0 | 1 | 1 | 0 | 1 | 0 | 1 | 1 | 1 | 1 | 1 | 1 | 1 | 0 | 1 | 1 | 1 | 1 | 1 | 1 | - | - | - | - | - | - | - | - | - | - | - | - | - |
| LB | 0 | 0 | 0 | 0 | 0 | 0 | 0 | 0 | 0 | 0 | 0 | 0 | 0 | 0 | 0 | 0 | 0 | 0 | 0 | 0 | 0 | 0 | - | - | - | - | - | - | - | - | - | - | - | - |
| MM | 0 | 0 | 1 | 1 | 0 | 1 | 0 | 3 | 1 | 1 | 1 | 1 | 1 | 1 | 0 | 3 | 2 | 1 | 1 | 1 | 1 | 0 | 3 | - | - | - | - | - | - | - | - | - | - | - |
| MED | 0 | 0 | 1 | 1 | 0 | 1 | 0 | 2 | 1 | 1 | 1 | 1 | 1 | 1 | 0 | 1 | 1 | 1 | 2 | 1 | 1 | 0 | 1 | 4 | - | - | - | - | - | - | - | - | - | - |
| NB | 0 | 0 | 0 | 0 | 0 | 0 | 0 | 0 | 0 | 0 | 0 | 0 | 0 | 0 | 0 | 0 | 0 | 0 | 0 | 0 | 0 | 0 | 0 | 0 | 1 | - | - | - | - | - | - | - | - | - |
| OV | 0 | 0 | 2 | 2 | 0 | 1 | 0 | 1 | 1 | 1 | 2 | 1 | 1 | 1 | 0 | 1 | 1 | 2 | 1 | 1 | 1 | 0 | 1 | 1 | 0 | 4 | - | - | - | - | - | - | - | - |
| PAAD | 0 | 0 | 1 | 1 | 0 | 1 | 0 | 3 | 1 | 1 | 1 | 1 | 1 | 1 | 0 | 2 | 2 | 1 | 1 | 1 | 1 | 0 | 2 | 1 | 0 | 1 | 4 | - | - | - | - | - | - | - |
| PRAD | 0 | 0 | 1 | 2 | 0 | 1 | 0 | 1 | 1 | 1 | 2 | 1 | 2 | 1 | 0 | 1 | 1 | 2 | 1 | 1 | 1 | 0 | 1 | 1 | 0 | 1 | 1 | 3 | - | - | - | - | - | - |
| RHAB | 0 | 0 | 0 | 0 | 0 | 0 | 0 | 0 | 0 | 0 | 0 | 0 | 0 | 0 | 0 | 0 | 0 | 0 | 0 | 0 | 0 | 0 | 0 | 0 | 0 | 0 | 0 | 0 | 0 | - | - | - | - | - |
| SKCM | 0 | 0 | 1 | 2 | 0 | 1 | 0 | 3 | 1 | 1 | 3 | 2 | 2 | 1 | 0 | 3 | 2 | 3 | 1 | 2 | 1 | 0 | 2 | 1 | 0 | 1 | 1 | 2 | 0 | 10 | - | - | - | - |
| STAD | 0 | 0 | 4 | 5 | 0 | 1 | 0 | 4 | 1 | 1 | 3 | 2 | 4 | 1 | 0 | 2 | 2 | 3 | 1 | 2 | 1 | 0 | 2 | 1 | 0 | 1 | 3 | 2 | 0 | 2 | 11 | - | - | - |
| STS | 0 | 0 | 0 | 0 | 0 | 0 | 0 | 0 | 0 | 0 | 0 | 0 | 0 | 0 | 0 | 0 | 0 | 0 | 0 | 0 | 0 | 0 | 0 | 0 | 0 | 0 | 0 | 0 | 0 | 0 | 0 | 0 | - | - |
| THCA | 0 | 0 | 0 | 0 | 0 | 0 | 0 | 2 | 0 | 0 | 0 | 1 | 0 | 0 | 0 | 1 | 0 | 1 | 0 | 0 | 0 | 0 | 1 | 0 | 0 | 0 | 0 | 0 | 0 | 2 | 0 | 0 | 3 | - |
| UCEC | 0 | 0 | 5 | 5 | 0 | 1 | 0 | 6 | 1 | 1 | 4 | 3 | 4 | 1 | 0 | 3 | 2 | 3 | 2 | 2 | 1 | 0 | 3 | 2 | 0 | 1 | 2 | 3 | 0 | 3 | 5 | 0 | 1 | 17 |

Notes: The full names of cancers are in Table S1.

Table S3. Uniquely significant genes of different cancers by WITER

| Cancer | Sig.Gene | UniqueGene | PubMedID |
| --- | --- | --- | --- |
| BLCA | 17 | 9 | CDKN1A[1.30e-16(30138622;29966976;29928692;29692692;29602637;29342159;29222807;28802642;28540176;28386678;27612592;27206339;26582573;24777035;24349619;23809295;23571005;23053941;18190825;11118045)], KDM6A[1.56e-12(30317582;29928692;29573965;29143738;29136510;28365159;28339163;28228601;27699256;27270441;26901314;26556859;25316812;25225064;24835989;24357085;23887298)], ZNF624[2.79e-11(?)], ELF3[1.33e-10(30528231;27612592;27699256;27217782)], ERCC2[5.12e-10(30290956;29980530;29603871;29133939;28803404;28802642;28707579;27479538;27376129;25636205;25316812;25096233;24504678;24121791;21864546;21426550;20061190;19706757;19429237;19242824;18815927;18630471;16537713)], RARG[4.28e-09(28078605)], FGFR3[1.76e-08(30344944;30342880;30258198;30220708;30213523;30171455;30154342;30128829;30064409;29941343;29876236;29791078;29673906;29654068;29573965;29525349;29517810;29487377;29463565;29435122;29423038;29396293;29262532;29226855;29193225;29165379;29161613;28927152;28855393;28775129;28760909;28601352;28597078;28589387;28583311;28548125;28523296;28521404;28507621;28416604;28388658;28365159;28320388;28255027;28247712;28229968;28169993;28108151;27932416;27930669;27846280;27835695;27530957;27612592;27726892;27699256;27669755;27611947;27596295;27586786;27523968;27455559;27447553;27381494;27356691;27352265;27268915;27197067;27189274;27157475;27029078;27029060;26924873;26901314;26869289;26861974;26778714;26722237;26715272;26651075;26598538;26547270;26542242;26522772;26452803;26374428;26351323;26327355;26278805;26274747;26254388;26244699;26238783;26171066;26144336;26041878;26039708;25997541;25993148;25809917)], RXRA[1.10e-07(30176882;29143738;28923856;26901314;26008846)], EP300[3.02e-06(30570744;29786110;28970362;27699256;27268145;26556859;22285928;21822268;18845648)] |
| BRCA | 19 | 8 | GATA3[8.47e-81(30554919;30468800;30389437;30354850;30332671;30327465;30271991;30251680;30246500;30185896;30134648;30125992;30125424;30100743;30062102;30061207;30035249;29997232;29972720;29902578;29894722;29854292;29792595;29662164;29609951;29593425;29546532;29535312;29510139;29462945;29435983;29431200;29416660;29408697;29358704;29351903;29262572;29207126;29202657;29123100;29053396;28966727;28965624;28945747;28884749;28810293;28805661;28789340;28752189;28722108;28703335;28693516;28690657;28611201;28581515;28580595;28574279;28514748;28428285;28423734;28394898;28351929;28288473;28273452;28258171;28211079;28078827;28077797;28066512;28038704;28027327;27997592;27917009;27904775;27900363;27867016;27829216;27809618;27666519;27654269;27588951;27556500;27556158;27514395;27473079;27356755;27354564;27338760;27283966;27184484;27154416;27093921;27041579;27018307;26998104;26960396;26922637;26907767;26852374;26825466)], MAP3K1[1.67e-29(30305115;30181556;30126855;30035249;29765551;29559730;29372690;29371908;29339359;29296238;29139094;28985766;28757652;28672935;28608266;28580595;28491135;28408616;28344865;28178648;28029147;28027327;27572905;26920143;26803517;26770289;26759750;26695891;26458823;26094658;25798844;25529635;24993294;24759887;24743323;24595411;24386504;24340245;24253898;24218030;24177593;23634849;23577780;23544012;23225170;23000897;22993404;22965832;22910930;22722202;22722201;22722193;22532573;22452962;21996731;21791674;21748294;21475998;21445572;21415360;21197568;21118973;20809358;20690207;20605201;20554749;19887619;19843670;19656774;19617217;19607694;19232126;19094228;19092773;19088016;19028704;18973230;18785201;18612136;18437204;18355772;17997823;17529967)], AKT1[2.68e-23(30537493;30445978;30443844;30443181;30420728;30394935;30343280;30337563;30289335;30285010;30212483;30185896;30154367;30140695;30126855;30065942;30035249;29963112;29893769;29844859;29808317;29783106;29769144;29658179;29636108;29589138;29550329;29540052;29506489;29482551;29365031;29202330;29103666;29090098;29086897;29074537;29053567;29018571;28973975;28964785;28945887;28903409;28853539;28806945;28693279;28685160;28569218;28495456;28478612;28476032;28446242;28423632;28356768;28301567;28287129;28116522;28061785;28036090;28027327;28008153;27956098;27917007;27806348;27769065;27699769;27748846;27738081;27672107;27634459;27628192;27608432;27600289;27574028;27531819;27515171;27374081;27302072;27297869;27197157;27095739;27059323;27035628;27028851;27004402;26926684;26899600;26741489;26710692;26700476;26631969;26608463;26549231;26540293;26536102;26500095;26474971;26400847;26351323;26302489;26267533)], CBFB[7.23e-17(28077088;28027327;26870154;26643573;22722202;22722193;9586906)], MAP2K4[3.90e-14(30563991;30035249;29854292;29765551;29755676;28491135;28446401;28344865;28027327;27792260;26907767;26249178;25086928;24194916;22722193;22522925;19593635;19404734;15578079;12097290;11754110)], TBX3[3.05e-08(30205045;29344954;28238063;28215225;27632063;27553211;27100732;26920143;26579496;26451490;26249178;26215676;25552398;25343378;23733266;23624936;22722201;22535523;22532574;22039763;21098263;20942798;21779450;19858224;19828084;19403417;19218121;18245468;18025091;16049973;15781639;15289316;11255752)], FOXA1[1.67e-06(30572598;30468461;30405052;30352905;30261643;30246500;30206966;30205045;30182384;30146936;30139998;30125992;30054444;29997232;29972720;29970021;29959737;29880907;29864144;29792595;29758547;29755131;29416660;29396764;29358704;29180470;29123100;28943920;28884749;28867731;28865492;28816236;28789340;28756535;28658208;28534958;28514748;28455227;28361702;28350011;28336670;28273452;28270510;28215225;27997592;27959926;27926873;27835577;27185372;27791031;27672107;27524420;27514395;27499099;27496708;27473079;27390128;27378691;27284343;27233940;27212698;27197147;27103403;27062924;27045898;27034986;27005559;26926684;26919034;26708273;26541755;26537518;26527523;26510790;26476779;26451490;26431101;26404658;26363213;26298189;26260807;26160249;26008846;25995231;25994056;25762479;25755696;25752574;25716347;25707489;25652398;25531315;25435372;25422910;25415051;25264199;25248036;25234841;25223786;25175082)], ZFP36L1[3.55e-06(29880481;19146866;17855657)] |
| KICH | 10 | 6 | VHL[5.90e-290(30581339;30565303;30555801;30553971;30523454;30522901;30502717;30513765;30372397;30365137;30355451;30349421;30340515;30291511;30289716;30277889;30173145;30149673;30141993;30076415;30072823;30066860;30026228;29938199;29872221;29805741;29732003;29720560;29674707;29662646;29550805;29527128;29503246;29481555;29479523;29463811;29342457;29245961;29239102;29218250;29215599;29202733;29194709;29176561;29169846;29144820;29141220;29118224;29100286;29097620;29095068;29033582;28925400;28893800;28853079;28812986;28779136;28765116;28731045;28723536;28701475;28667082;28632992;28624320;28582447;28553932;28550387;28543713;28533271;28488172;28473526;28408295;28329682;28261513;28259286;28257806;28256713;28247252;28235946;28120493;28052007;27923499;27909050;27845047;27841867;27836247;27595394;27774982;27764136;27751729;27741516;27728802;27682873;27623354;27556922;27530247;27506904;27491826;27491085;27485825)], PBRM1[4.70e-90(30446457;30372397;30355451;30349421;30277889;30072823;30033103;30009957;29674707;29617669;29426696;29301960;29218250;29169846;29158875;29095068;28921948;28812986;28779136;28765116;28731045;28723536;28618948;28473526;28459210;28445125;28408295;28329682;28327121;28284891;28212566;28092369;28053089;27764136;27751729;27556922;27407122;27100670;26891804;26864202;26484545;26452128;26300218;26166446;26111976;26003625;25997916;25873528;25676555;25628030;25583177;28326264;25465300;25124064;24992170;24821879;24186201;24166983;24158655;24053427;24029645;23792563;23620406;23416164;23036577;22949125;22805307;22461374)], BAP1[3.30e-37(30519310;30446457;30405850;30372397;30355451;30349421;30277889;30269473;30135306;30072823;29981911;29853346;29617669;29558292;29426696;29266978;29158875;29118224;28900502;28812986;28779136;28765116;28753773;28731045;28723536;28618948;28488170;28473526;28459210;28408295;28327121;28284891;28212566;27751729;27556922;27085487;26891804;26864202;26854086;26839909;26484545;26452128;26300492;26300218;26166446;26111976;25972334;25873528;25826081;28326264;25479927;25465300;25126716;25124064;24821879;24382589;24166983;24158655;24128712;24076305;24029645;23709298;23620406;23277170;23036577;22949125;22805307;22461374)], SETD2[3.53e-24(30487242;30406665;30372397;30355451;30349421;30072823;30033103;29674707;29558292;28812986;28779136;28754676;28753773;28731045;28723536;28445125;28408295;28260718;27764136;27751729;27288695;27556922;27292023;26891804;26864202;26575290;26559293;26537074;26452128;26166446;26111976;26073078;25873528;25853938;25714014;28326264;25124064;24821879;24186201;24166983;24158655;24029645;23792563;23620406;23036577;22949125;22805307;22461374;20501857;20054297)], KDM5C[9.40e-22(30355451;30033103;28779136;28723536;28408295;28212566;27751729;27556922;26484545;25124064;24029645;23036577;22949125;21725364;20054297)], MTOR[6.10e-10(30581339;30524898;30513765;30488209;30368665;30349421;30072823;30016766;29610387;29536301;29508215;29479523;29202733;29158991;29144820;29118224;29079709;28927098;28779136;28765116;28723536;28680592;28545465;28396841;28329682;28257806;28247252;28197812;27751729;27738339;27615548;27574806;27453294;27405474;26974204;26787754;26609489;26408740;26255626;26224474;25997916;25948777;25783986;25625928;25520878;25426415;25351205;25293974;25186283;25152703;24929890;24821879;24575852;24565854;24504440;24496460;24495452;24136229;23797736;23633458;23290145;22322364;21798997;21644050;20709527;20437403;20022054;19956876;19843858;19663736;19657325;19265534;17987219)] |
| UCEC | 17 | 5 | FGFR2[2.17e-15(30094104;30002137;28953502;28119489;27550940;26626801;26366417;26294741;25794154;25720322;25517871;25019571;24696723;24448819;23778141;23536011;23300780;22383975;21725289;21372219;21221135;19147536;19078924;18785201;18636142;18552176)], PPP2R1A[7.24e-13(30104481;29915797;28940304;28718916;28485815;28292439;27499902;27485451;27272709;26626801;25720322;25308272;23588898;23359684;22653804)], CHD4[2.29e-10(29888111;29844320;28940304;27997699;23359684;23104009)], CCND1[5.35e-07(30585737;30482501;29969496;29232554;28408839;27831653;27648123;27349856;27105504;26366417;26353976;25546926;25214561;24779718;24337234;24126431;23733133;23731275;21454826;16569247;15069681;12955092;10473073)], FOXA2[1.89e-06(30091462;29546371;29442045;28940304;27538367;25994056;22945641)] |
| HNSC | 12 | 5 | NOTCH1[3.51e-09(30087145;29970484;29909892;29771197;29747488;29489439;29340043;29331751;29232766;29146722;29068587;29053175;28195818;27965308;27595504;27380877;27117272;27035284;27028310;26927514;29034103;25836654;25633867;25588898;25580884;28324520;25440877;25303977;25275298;25234595;24787294;24670651;24667986;24292195;24277457;24001612;23750501;23714515;23645351;23607916;22773520;21798897;21798893;20175927;20127005;19550121)], AJUBA[2.68e-08(29771197;29053175;28126323;29034103;25303977)], EPHA2[3.18e-08(24864260;22455776;21955398;18425361)], NSD1[3.90e-07(29884412;29636367;29340043;29213088;29053175)], ZNF750[4.76e-07(26949921)] |
| COAD | 12 | 4 | APC[1.69e-109(30585891;30582224;30577807;30575318;30572883;30554113;30550822;30525086;30523670;30530050;30521203;30513515;30511962;30498084;30496442;30475060;30472235;30462522;30460349;30459318;30448068;30447686;30414835;30413483;30405780;30396459;30377564;30377189;30374053;30365932;30361697;30355737;30350313;30337690;30324682;30316137;30315134;30311185;30292635;30273442;30272267;30271205;30261643;30259713;30256826;30256815;30254720;30239619;30238564;30231850;30229128;30221070;30202242;30186819;30177706;30166531;30156908;30152102;30150766;30145695;30140377;30131849;30121899;30116794;30110600;30109253;30107178;30097126;30097032;30087854;30073936;30072583;30066856;30065793;30065304;30063922;30032163;30029640;30024920;30001952;29980571;29976257;29972397;29967336;29967250;29964340;29955725;29945573;29937994;29931584;29914973;29901124;29895971;29891445;29880585;29872723;29871936;29844865;29800151;29786110)], AMER1[2.05e-10(30062471;30032163;28551381;28002797;26527806;26071483;24251807)], SMAD2[1.35e-08(30536375;30476540;30412716;30182733;29973939;29967250;29945573;29844309;29690747;29599290;29545874;29453410;29449642;29443735;29219668;29050211;29037859;28833571;28714374;28601657;28592228;28551381;28445620;28299321;28243320;28106826;28049506;27856280;27840994;27662660;27528036;27401208;27330076;27222126;27089389;26647993;26370966;26328044;25980495;25974029;25937570;25934251;25889203;25736321;25653118;25422078;25238234;25169976;25166914;24970681;24811787;24727557;24696849;24668684;24627270;24427302;23886856;23867993;23661436;23536895;23426176;23153552;23139211;23108401;22452939;22307228;22227581;21998011;21978709;21747928;21702981;21480389;21473864;21365634;21068203;20732443;20622003;20432436;20126641;19798702;19686287;19571605;19147584;18949743;18781153;18769113;18425817;17893910;16828225;16288847;16015041;15963949;15711891;15520171;15374947;14720321;14715079;14520001;12967141;12866583)], TCF7L2[2.59e-08(30350386;30200414;30131849;30032163;30026326;29975781;29802748;29511559;29389519;29301589;29245969;29135090;29131639;29118424;29050326;28949031;28811361;28472810;28450117;28362475;28343235;28143522;28117551;28060743;28002797;27792933;27761963;27504909;27755946;27709738;27527215;27398792;26474385;26243311;26191083;26165840;26060019;25913757;25277775;25205133;25131200;25050608;24943349;24913975;24836286;24828199;24670930;24608966;24398765;24338422;24317174;23951231;23817222;23796952;23319804;22895193;22419714;22108803;22005519;21983179;21956205;21892161;20056134;19924301;19760027;19561607;18992263;18621708;18478343;18398040;18268068;18268006)] |
| LAML | 10 | 4 | DNMT3A[6.32e-25(30583461;30582132;30553776;30504903;30409182;30305534;30282643;30275526;30245403;30122995;30122013;30017658;29988143;29903761;29877252;29806051;29780592;29727682;29721667;29721207;29702001;29674693;29643943;29627222;29624746;29619119;29573577;29556023;29549529;29518238;29491461;29472724;29464054;29343972;29330206;29324392;29320732;29309772;29306105;29296935;29285580;29274134;29254789;29254227;29249818;29227476;29193057;29188605;29172276;29166738;29137279;29079128;29075615;29069784;29024628;28992762;28978861;28978821;28960408;28940816;28923882;28905428;28877686;28872462;28830460;28823257;28767575;28753595;28751771;28713819;28710806;28643785;28642303;28484171;28461508;28452374;28449304;28418922;28408400;28386358;28341738;28315400;28297624;28255022;28215704;28152414;28070990;28061354;28052028;28003281;27983727;27881874;27841873;27821287;27435003;27795554;27636548;27724883;27636998;27626217)], FLT3[1.85e-17(30568173;30555035;30555023;30553002;30552988;30544932;30501709;30514174;30510164;30500073;30472492;30472087;30466756;30466744;30441973;30428571;30420889;30410824;30410361;30405611;30389660;30378313;30372617;30369187;30357752;30344940;30341082;30332834;30320942;30305502;30303964;30301399;30275526;30266802;30250637;30245083;30190467;30190345;30190321;30180463;30148617;30146162;30122013;30117185;30111391;30102945;30089730;30082225;30081867;30064973;30046393;30017192;30015632;29983874;29959200;29951132;29907544;29903761;29894944;29857559;29806051;29784639;29773641;29773601;29773106;29766829;29748445;29721667;29716633;29696374;29692343;29688850;29682194;29665898;29664232;29663558;29654398;29654265;29643943;29625580;29624746;29573577;29563537;29556023;29551027;29541391;29534404;29530994;29507660;29505696;29491461;29487059;29472722;29472720;29472718;29463564;29463558;29437468;29431743;29416774)], NPM1[1.80e-07(30577887;30555023;30466744;30428571;30420649;30410824;30404199;30389521;30383325;30341082;30305534;30303964;30298506;30214626;30205049;30190467;30190321;30176240;30126426;30122995;30122013;30111391;30089730;30082225;30017658;30015632;29932212;29903761;29857559;29806051;29766829;29764005;29721667;29696374;29665898;29661468;29625580;29624746;29622865;29573577;29563537;29556023;29541391;29534404;29530994;29519869;29491461;29472722;29441887;29435155;29423110;29408852;29402726;29343273;29330746;29310020;29286103;29283500;29274134;29254789;29254227;29249819;29238371;29224316;29221119;29219176;29193057;29188605;29172276;29166740;29166738;29157973;29111347;29090521;29079128;29069784;28978861;28971903;28923882;28920929;28882949;28841206;28836868;28835438;28830460;28823257;28753595;28740552;28710806;28698788;28679652;28618016;28574487;28569789;28475434;28473620;28471807;28456748;28452374;28411256)], CEBPA[1.53e-06(30476680;30466750;30420667;30337300;30305534;30303964;30190467;30122013;30078804;30032569;30015632;29932212;29861846;29806051;29766829;29624746;29622865;29573577;29541391;29534404;29515250;29483711;29435155;29431622;29402726;29343483;29310020;29306105;29286103;29238371;29193057;29188605;29032147;29025912;28978861;28923882;28900037;28895127;28882949;28830460;28753595;28745571;28663557;28504718;28473620;28452374;28380436;28357685;28341738;28299657;28250006;28249600;28210006;28203345;28186500;28179278;28144729;28090023;28074068;28070990;27899775;27812248;27694926;27626217;27512765;27367478;27359055;27350755;27288520;27285584;27129260;27062340;27040395;27034432;27023522;27012040;26992835;26940274;26876264;26802049;26725349;26721895;26708912;26693794;26676635;26586702;26537612;26496024;26488113;26466372;26460249;26450903;26419342;26408402;26386075;26377688;26376842;26375248;26374622;26239249)] |
| SKCM | 10 | 4 | RAC1[2.70e-12(30509087;30197759;30157875;30110134;29673588;29600692;29491429;29432733;29342889;29222169;29095823;29059171;28915620;28803992;28562340;28416823;28277539;28210865;27908735;27890785;27846383;27715393;27575453;27699663;28947953;27418645;27314100;27181209;27121131;27111337;26884185;26782071;26744134;26392417;26343386;26341689;26248315;26202910;26176707;26079945;25857817;25677173;25505176;25465943;25452114;25203554;25056119;25043693;24755198;24690323;24659802;24659799;24625986;24586549;24531394;24514042;24480879;26263704;24468268;24441506;24387669;24358134;24341237;24337068;24265417;24200678;24088985;23981010;23957481;23382862;23372575;23337888;23330781;23284172;23222813;22940494;22843693;22842228;22817889;22718121;22469839;22441674;22335598;22174906;21765460;21698524;20869211;20803552;20631911;20236250;19881949;19276388;18835169;18815275;18728402;18577517;18495474;18425380;18089842;17952876)], MAP2K1[6.39e-09(30201825;29461977;28881731;26913480;26684394;26673799;26343386;26018731;23639941;23444215;23174022;22197931;22105811;21726664;20526349)], PPP6C[1.07e-08(30509087;26868000;25857817;25486434;24755198;26263704;24341237;24336958;23729733;22842228;22817889)], DSG3[3.54e-06(11422052)] |
| LUAD | 8 | 4 | STK11[1.02e-23(30297789;30297358;30279957;30275180;30224757;30075702;30050779;30031117;29898990;29787863;29773717;29764856;29575851;29540834;29535211;29337640;29307989;29279706;29219616;29198084;29191602;29168346;29066508;28914263;28911955;28884744;28754670;28652249;28619094;28538732;28435024;28413430;28387316;28336552;28205554;28145643;27923066;27687306;27565922;27467949;27299180;27218826;27151654;27121209;26960398;26917230;26833127;26829311;26625312;26599269;26477306;26463840;26420428;26350096;26124082;26119936;26087898;26069186;26066407;25982285;25969368;25964588;25695224;25634010;25477232;25444907;25278450;25122068;25036236;25031567;24828662;24482041;24468202;24448687;24297535;24236184;24086281;24077454;24054548;23276293;23047306;22768234;22590557;21532627;20057966;19661141;19483050;19353596;19176640;19165201;17711506;17676035;17216011;16912160;16580634;15639728;15021901;12097271;11212897;10508479)], U2AF1[3.87e-09(29991672;27776121;27545006;24498085;22980975;9490301)], EGFR[2.50e-07(30586539;30584328;30584321;30582673;30580903;30580372;30578931;30578392;30576239;30572543;30572427;30568033;30565869;30560887;30555679;30555633;30555330;30553753;30546903;30546439;30544036;30543838;30524878;30524640;30523199;30522449;30522171;30522170;30522169;30540703;30537515;30536734;30536070;30535340;30529597;30527195;30526458;30521971;30518123;30508797;30505713;30505499;30505497;30516345;30515095;30513627;30500458;30499112;30483773;30483119;30480362;30475455;30475204;30473379;30470932;30470824;30463991;30461012;30454551;30454542;30453282;30452286;30451486;30445769;30445271;30442274;30431083;30429415;30429043;30429037;30429033;30429031;30429024;30425968;30425905;30420490;30419315;30417422;30413663;30412601;30408128;30406144;30406002;30404194;30401947;30400936;30395779;30393484;30391576;30391211;30390416;30390072;30389917;30389658;30382076;30381940;30379408;30378264;30368879;30368012)], RIT1[2.98e-06(30391781;29636358;26098749;25079552;24469055)] |
| LGG | 7 | 3 | ATRX[1.51e-12(30194745;29111096;29091765;29027701;28980701;28419269;28392842;27758882;26508407;25664944;24710217;23373454)], FUBP1[3.40e-07(29606613;23373454)], CIC[9.23e-07(30093628;23373454)] |
| STS | 11 | 2 | RHOA[6.54e-11(29167336;28709411;28196303;27715394;26819309;26696895;28174704;26480288;25904553;25838983;25793574;25610737;25569678;25471072;25253349;23645742;23580575;22696654;18521231;17881360;17507466;17005646;16448388;12847101;11590141;1710770)], TLR4[3.75e-06(29051178;28421060;25320320;24899179;23723066;23514335;23365457;18996347)] |
| MED | 4 | 2 | DDX3X[4.64e-21(29582169;29222110;27180681;27058758;26290144;25724843;24608801;22832583;22820256;22722829)], SMO[3.24e-06(30554998;30487124;30483764;30452905;30374857;30106753;29857275;29777201;29531057;29378965;29348431;29274272;29208776;29055107;28923910;28873303;28833911;28716052;28618224;28605510;28487292;27785591;27495899;27236920;27069629;26960983;26891329;26691947;26633513;26450969;26371509;26323341;26286140;26169613;26113054;26080084;25859932;25636740;25505589;25485584;25484239;25376612;25355313;25306392;25131638;24994715;24973920;24951114;24871706;24276242;24068730;23872071;23671675;23662017;22966790;22923130;22869526;22851551;22452947;24451804;22084163;21618411;21501498;21325292;21143927;21123452;20881279;20524040;20493695;20386868;20024066;19726788;19701203;18826648;18502968;18288402;17413002;17017853;16707575;16618744;15806168;12192414;11965540;10984056;10564585)] |
| OV | 4 | 2 | BRCA1[3.72e-10(30584090;30576872;30570007;30561604;30555256;30553462;30552672;30541753;30524945;30538879;30535581;30506513;30517521;30498870;30482293;30480775;30473125;30472649;30471648;30458859;30453575;30441849;30426508;30414230;30410611;30400234;30392916;30384980;30383754;30382883;30364812;30358186;30353044;30352249;30348217;30342021;30333958;30333088;30327455;30325501;30322717;30315757;30309222;30309218;30308700;30306255;30303537;30295091;30283497;30275862;30275608;30268633;30263092;30262796;30254663;30235698;30233647;30228165;30221688;30219179;30215333;30209399;30209015;30207098;30191368;30186769;30186165;30141832;30132561;30128536;30116283;30111881;30111871;30110192;30105462;30103829;30101128;30100923;30092674;30078507;30075112;30055521;30050740;30049288;30045136;30042414;30042185;30040829;30036195;30018258;30014164;29998407;29998185;29997359;29996917;29988077;29982601;29975922;29969168;29961768)], CDK12[1.10e-07(30422115;30333958;30104286;29872499;28950147;27905519;27662623;27241520;26247403;25429106;24554720;24240700;22012619;21720365)] |
| PAAD | 4 | 1 | TGFBR2[1.48e-07(29899418;29393426;28809762;28373289;26279302;26255562;25791160;23690952;23378339;23237571;23103869;22523087;17031113;12615714;9850059;9598801)] |
| DLBCL | 2 | 1 | GNA13[2.29e-07(28302137;27980305;26989201;26819451;26773040;26616858;26608593;25991819;25274307;23699601;23292937;23143597;22343534)] |
| NB | 1 | 1 | ALK[6.59e-07(30538293;30459283;30459281;30425456;30400214;30375995;30350464;30266251;30117275;30115695;30061385;30013190;29891519;29753117;29660984;29642598;29638111;29600072;29559559;29556564;29555900;29535836;29515255;29505958;29492199;29466695;29455642;29441070;29380702;29378002;29374774;29371588;29357780;29321660;29317532;29296183;29290991;29203817;29184034;29084134;29081033;29069774;29027209;29018329;28915622;28915608;28871274;28800395;28756644;28676342;28674118;28666189;28665006;28662353;28604107;28602975;28546523;28521285;28458126;28425916;28423360;28350380;28338501;28326957;28178969;28163672;28139105;28069802;28030793;27997549;27879258;27830764;27707976;27684973;27655666;27604320;27573755;27483357;27471553;27285993;27179218;27165366;27076624;27013922;27009859;27009842;26986945;26925973;26893860;26835380;26829053;26826611;26794043;26750252;26735175;26687816;26633716;26630010;26616860;26539795)] |
| LUSE | 8 | 0 | - |
| GBM | 6 | 0 | - |
| MM | 3 | 0 | - |
| PRAD | 3 | 0 | - |
| THCA | 3 | 0 | - |
| CLL | 2 | 0 | - |
| LIHC | 2 | 0 | - |
| ESCA | 1 | 0 | - |
| KIRC | 1 | 0 | - |
| LUSC | 1 | 0 | - |
| ALL | 0 | 0 | - |
| AT | 0 | 0 | - |
| CARC | 0 | 0 | - |
| CESC | 0 | 0 | - |
| KIRP | 0 | 0 | - |
| LB | 0 | 0 | - |
| RHAB | 0 | 0 | - |
| STAD | 0 | 0 | - |

**Note**: The values in square brackets are p-values by WITER. The significant genes are determined according to Bonferroni correction for family-wise error rate (<0.05). The number in the brackets are the PubMed ID of papers co-mentioning the disease name and gene symbol according to search API in PubMed database, [http://eutils.ncbi.nlm.nih.gov/entrez/eutils/esearch.fcgi?db=pubmed&term=“DiseaseNames(inlcuding](http://eutils.ncbi.nlm.nih.gov/entrez/eutils/esearch.fcgi?db=pubmed&term=) homonymies)”[tiab]%29+AND+“GeneSymbol (including RefSeq mRNA IDs)” [tiab]. For genes with over 100 papers, only the most recent 100 papers are shown. “-” denotes no unique significant genes. “a”: number of total significant genes. “b”: number of unique significant genes. The full names of cancers are in Table S1.

Table S4: Rescued genes in the Breast invasive carcinoma dataset by different tools

|  |  | SubSample | Missed | Rescued by | | | |
| --- | --- | --- | --- | --- | --- | --- | --- |
| Tested tool^a^ | **RandomSampleID** | **Sig.Genes** | **Sig.Genes** | **MutSigCV** | **oncodriveFML** | **WITER** | **20/20plus** |
| MutSigCV (16) | 1 | 10 | 6 | -c | 0 | 2 | 2 |
|  | 2 | 8 | 8 | - | 0 | 3 | 2 |
|  | 3 | 10 | 6 | - | 0 | 3 | 2 |
|  | 4 | 7 | 9 | - | 1 | 3 | 3 |
|  | 5 | 8 | 8 | - | 2 | 4 | 1 |
|  | 6 | 8 | 8 | - | 0 | 4 | 4 |
| OncodriveFML (9) | 1 | 5 | 4 | 0 | - | 0 | 1 |
|  | 2 | 5 | 4 | 0 | - | 2 | 1 |
|  | 3 | 6 | 3 | 0 | - | 1 | 1 |
|  | 4 | 6 | 3 | 0 | - | 1 | 1 |
|  | 5 | 5 | 4 | 0 | - | 1 | 1 |
|  | 6 | 6 | 3 | 0 | - | 2 | 2 |
| WITER (19) | 1 | 14 | 5 | 0 | 0 | - | 0 |
|  | 2 | 16 | 3 | 0 | 0 | - | 0 |
|  | 3 | 14 | 5 | 0 | 1 | - | 1 |
|  | 4 | 14 | 5 | 0 | 0 | - | 0 |
|  | 5 | 14 | 5 | 0 | 0 | - | 0 |
|  | 6 | 17 | 2 | 0 | 0 | - | 0 |
| 20/20plus (18) | 1 | 11 | 7 | 1 | 1 | 1 | - |
|  | 2 | 8 | 10 | 0 | 0 | 4 | - |
|  | 3 | 11 | 7 | 1 | 0 | 3 | - |
|  | 4 | 8 | 10 | 0 | 0 | 1 | - |
|  | 5 | 9 | 9 | 1 | 1 | 3 | - |
|  | 6 | 11 | 7 | 0 | 0 | 1 | - |

a: The significant genes detected by a tool in the full dataset. b:The significant genes detected by a tool in the extracted random samples. c: This item is inapplicable.
